# Arp2/3-mediated bidirectional actin assembly by SPIN90 dimers in metazoans

**DOI:** 10.1101/2025.01.31.635869

**Authors:** Tianyang Liu, Luyan Cao, Miroslav Mladenov, Guillaume Romet-Lemonne, Michael Way, Carolyn A. Moores

## Abstract

Branched actin networks nucleated by the Arp2/3 complex play critical roles in various cellular processes, from cell migration to intracellular transport. However, when activated by WISH/DIP/SPIN90 family proteins, Arp2/3 nucleates linear actin filaments. Unexpectedly, we found that human SPIN90 is a dimer that can nucleate bidirectional actin filaments. To understand the basis for this, we determined a 3 Å resolution structure of human SPIN90-Arp2/3 complex nucleating actin filaments. Our structure shows that SPIN90 dimerises via a 3-helix bundle and interacts with two Arp2/3 complexes. Each SPIN90 molecule binds both Arp2/3 complexes to promote their activation. Our analysis demonstrates that single filament nucleation by Arp2/3 is mechanistically more like branch formation than previously appreciated. The dimerisation domain in SPIN90 orthologues is conserved in metazoans, suggesting that this mode of bidirectional nucleation is a common strategy to generate anti-parallel actin filaments.

## INTRODUCTION

The actin cytoskeleton is a versatile dynamic assembly of filaments formed by the polymerisation of actin monomers that participates in many cellular processes. A major feature of the actin cytoskeleton is branched actin networks, which generate the forces necessary for membrane deformation, intracellular trafficking and cell migration^1–5^. The Arp2/3 complex, consisting of seven subunits (Arp2, Arp3 and ArpC1-C5), is the only known cellular constituent that can nucleate daughter actin filaments from the side of a pre-existing mother filament^1, 6–12^. In the presence of pre-existing mother filaments, activation of Arp2/3 by Class 1 nucleation-promoting factors (NPFs) induces a set of conformational changes within the complex that enable actin nucleation and thereby allow branch assembly^13–26^. These include twisting of the hinge helices in ArpC2 and ArpC4 that moves the actin-like subunit Arp2 towards Arp3 to produce an F-actin-like, short-pitch arrangement, with Arp2 and Arp3 each adopting a flattened structure^20, 22, 23, 25, 26^. Together these conformational changes create a template for the addition of actin monomers and the nucleation of a daughter filament (*Extended Data Fig. 1a*). While recent advances have clarified multiple aspects of the mechanism of branch formation, a central question remains: how is the initial mother filament formed? Since in cells actin monomers are in complex with profilin, spontaneous nucleation of actin filaments is rare^27^. This points to the crucial role of actin filament nucleators in defining the timing and orientation of linear mother filament formation prior to initiation of the branched actin network.

In yeast, the actin regulator Dip1 can activate Arp2/3 without the need for preformed mother actin filaments^28^. Arp2/3 mediated actin patch formation and endocytosis in yeast are delayed in the absence of Dip1, highlighting its importance in generating initial mother filaments for branch network formation^29, 30^. Dip1 is a member of the WISH/DIP/SPIN90 family, characterised by a conserved leucine-rich domain (LRD) that binds Arp2/3 (*Extended Data Fig.1b*)^28^. In mammals, the SPIN90 LRD sits within a C-terminal armadillo repeat domain, while N-terminal SH3 and poly-proline regions are involved, respectively, in SPIN90 localisation and interactions with the signalling adaptor Nck^31^. Importantly, however, residues in the region that connect the poly-proline region and the armadillo repeat domain (residues 274 – 376) are essential for activating the mammalian Arp2/3 complex ^28, 31, 32^ (Fig. 1a). The knock-down of mammalian SPIN90 leads to a loss of growth factor-induced lamellipodia formation and EGFR-mediated endocytosis^33, 34^. Moreover, SPIN90 depletion reduces cortical actin mesh size in blebs and stiffens the mitotic cortex, further highlighting its importance in regulating actin organisation and dynamics^35^. Together, these findings suggest that SPIN90 also seeds branched actin networks in mammalian cells by generating mother filaments.

**Figure 1.**
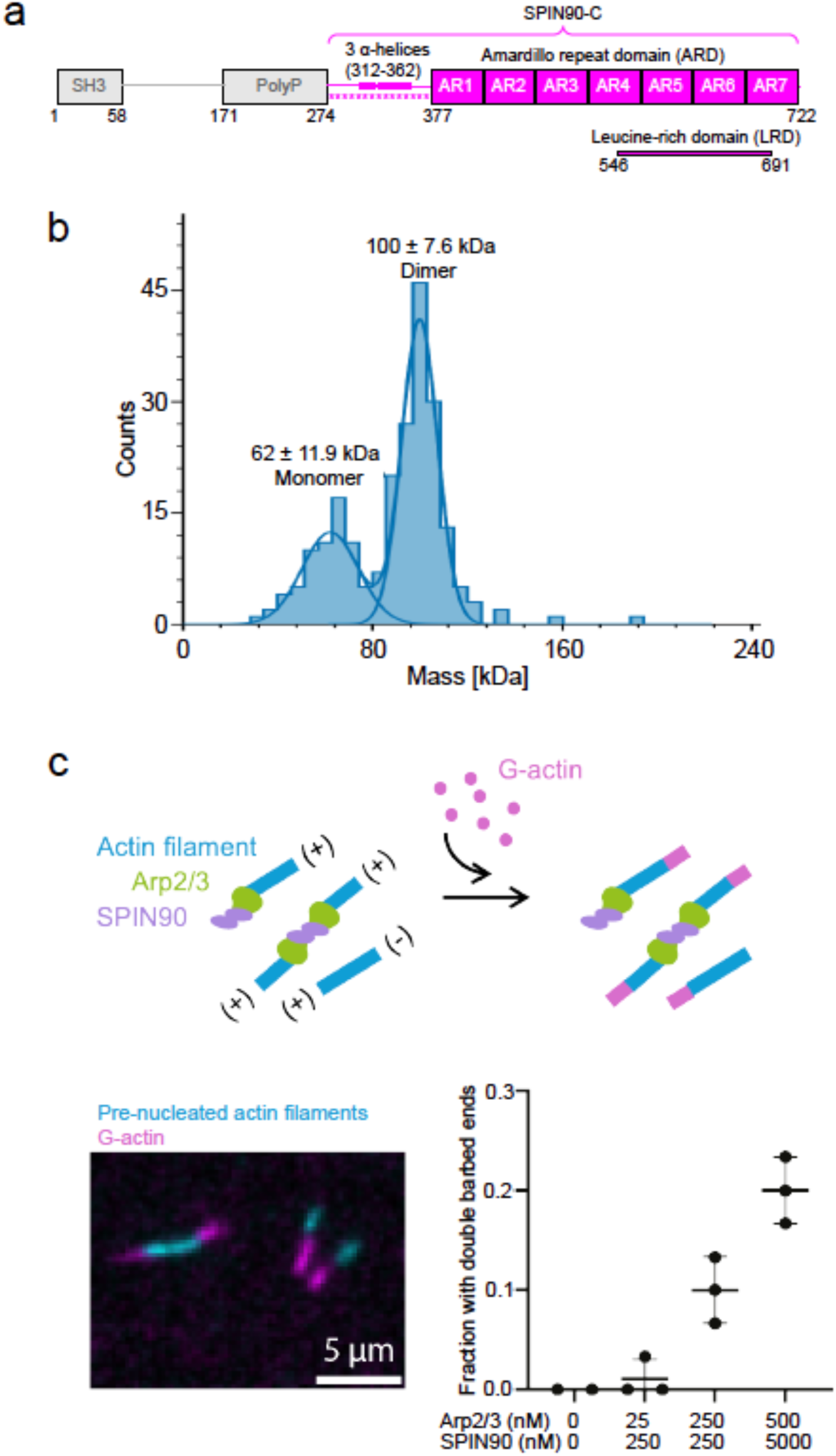
SPIN90 forms a dimer that induces bidirectional Arp2/3 nucleated actin filament polymerisation. a. Human SPIN90 (Q9NZQ3) domain organisation. The SPIN90-C construct used in this study is coloured in magenta. The N-terminal region of SPIN90-C that is essential for activating the Arp2/3 complex in addition to the armadillo repeat domain (ARD) is indicated with magenta stars^32^. b. Mass distribution of 30 nM SPIN90-C, whose theoretical molecular weight is 50.0 kDa. Two peaks were observed with molecular weight corresponding to 100 ± 7.6 kDa and 62 ± 11.9 kDa, showing that SPIN90-C exists mainly as a dimer in solution. c. Schematic (top) of in vitro TIRF microscopy assay to investigate assembly of actin filaments from two ends of SPIN90 activated Arp2/3 complex. Pre-polymerised actin filaments (15% labelled with Alexa-568, cyan) were mixed with G-actin (15% labelled with Alexa-488, magenta). The barbed ends were visualised directly by the Alexa-488 signal (bottom left). Quantification of fraction of actin filaments with double barbed ends (bottom right). Each point represents the result of an independent experiment. The bars show the mean and standard error.

The cryo-EM structure of Dip1-activated yeast Arp2/3 at the end of a nucleated actin filament reveals that the LRD of Dip1 binds to the ArpC4 hinge helix (*Extended Data Fig. 1b*) ^36^. The same interaction is also observed in the co-crystal structure of human SPIN90 armadillo repeat domain bound to inactive bovine Arp2/3^32^. It was therefore proposed that WISH/DIP/SPIN90 family proteins activate Arp2/3 by a conserved mechanism that involves interacting with and bending the ArpC4 hinge helix. This in turn promotes the Arp2-Arp3 short-pitch conformation necessary to nucleate an actin filament. Although current structural insights have deepened our understanding of how WISH/DIP/SPIN90 family proteins interact with and activate the Arp2/3 complex, several questions remain. In particular, how does the small interface between WISH/DIP/SPIN90 proteins and ArpC4 fully activate Arp2/3, especially when activation during actin branch formation requires extensive interactions with the mother filament (*Extended Data Fig. 1a, b*)? Furthermore, what contribution do residues 274 – 376 N-terminal to the armadillo repeat domain of mammalian SPIN90 make to Arp2/3 activation? To address these questions, we examined the interaction of human SPIN90 with human Arp2/3 complex using biophysical and structural approaches. We found that SPIN90 forms a dimer via a 3-helix domain N-terminal to the armadillo repeats, facilitating bidirectional actin filament nucleation by activating two Arp2/3 complexes. The 3-helix domain and predicted dimeric structure are present in multicellular animals, suggesting that this mechanism of bidirectional actin filament nucleation is conserved in metazoans.

## RESULTS

### SPIN90 dimers induce bidirectional Arp2/3 nucleated actin filament polymerisation

To better understand how SPIN90 promotes Arp2/3-mediated actin nucleation, we first studied the biophysical properties of the minimal functional region of recombinant human SPIN90, which lacks its signalling and localisation domains (SPIN90-C, Fig. 1a). Using mass photometry we observed a major peak with a molecular mass of 100 kDa, approximately corresponding to the size of a SPIN90-C dimer, and a smaller monomeric peak (Fig.1b). To explore the implications of this unexpected SPIN90 dimerisation for Arp2/3 activation, we performed TIRF microscopy (Fig. 1c). We first polymerised short, fluorescently-labelled actin seeds (labelled with Alexa-568, cyan) in the presence of SPIN90 and the Arp2/3 complex, then added actin monomers (labelled with Alexa-488, magenta) and visualised dynamic filament growth. Whereas activated Arp2/3 usually nucleates unidirectional actin polymerisation, we found that with increasing concentrations of SPIN90 together with Arp2/3, more actin polymerised from both ends of the actin seeds (Fig. 1c). This demonstrates that SPIN90 dimerisation can drive bidirectional Arp2/3-mediated actin filament growth.

To understand the mechanism of SPIN90 dimerisation and its contribution to actin nucleation, we used cryo-electron microscopy (cryo-EM) to determine the structure of the SPIN90-Arp2/3 complex in the presence of actin (*Extended Data Fig. 1*). The resulting reconstruction (at approximately 3 Å overall resolution (*Extended Data Fig. 2-3*) revealed a SPIN90 C2-symmetric dimer at its centre, flanked by two activated Arp2/3 complexes nucleating actin filaments in a bidirectional fashion (Fig. 2a, Movie. 1). 3D variability analysis of the C1 structure reveals some stretching due to intrinsic flexibility of the SPIN90 armadillo repeats (Movie. 2), a well-known property of these extended, curved domains^37^. Our structure shows that the SPIN90 Dimerisation Domain (SDD) is formed from three α-helices (residues 312-362) that interact through extensive rigid, hydrophobic contacts (Fig. 2b, c, Movie.1). The SDD is connected by a flexible loop (residue 363-376) to the seven armadillo repeats (residues 377-717) (Fig. 1a, 2b), while the N-terminal 38 residues of SPIN90-C are not visible, likely due to flexibility.

**Figure 2.**
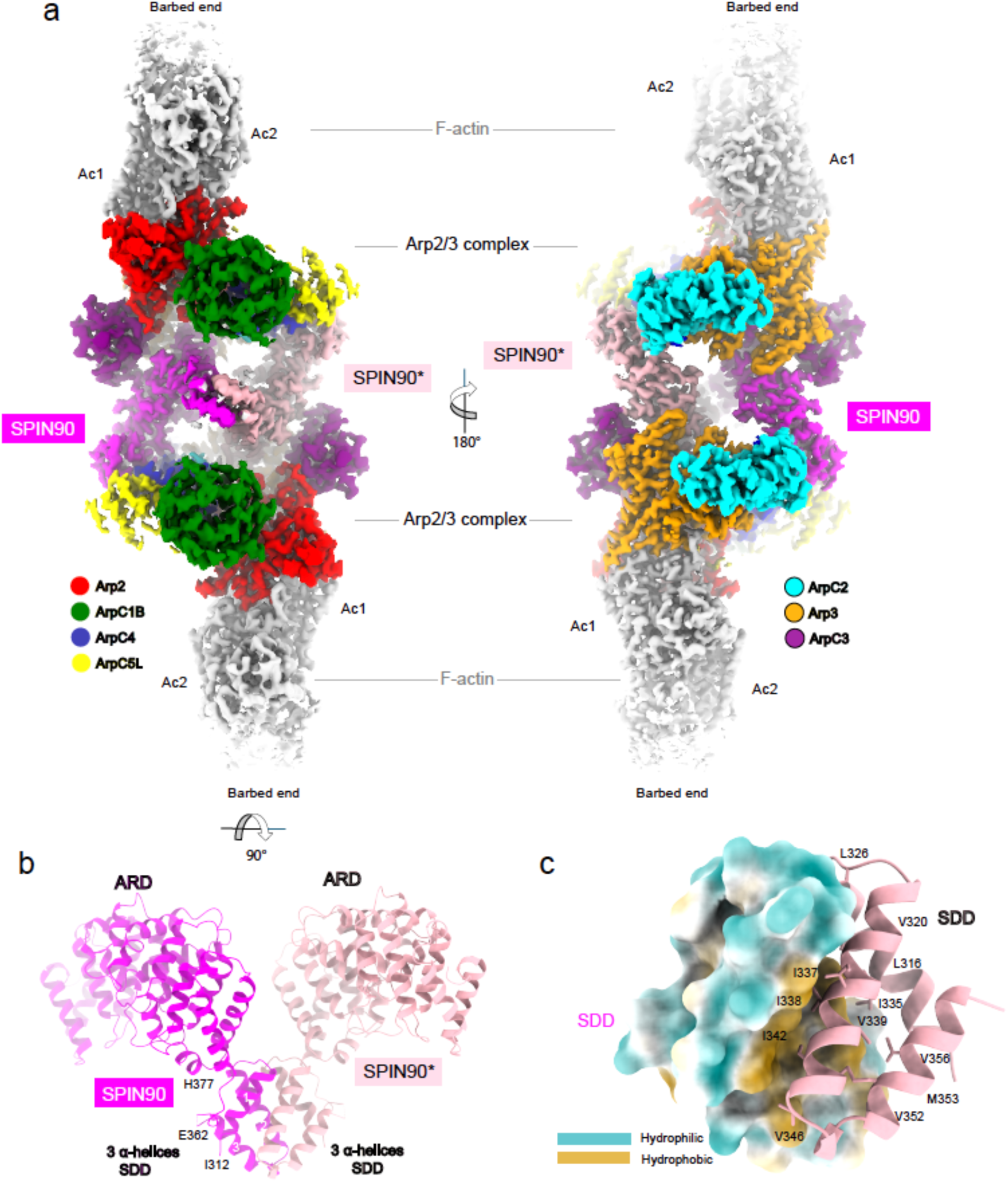
Structure of the SPIN90-Arp2/3 nucleated bidirectional actin filaments. a. Overview of the cryo-EM reconstruction of the SPIN90-Arp2/3 complex nucleated bidirectional actin filaments. Densities of individual proteins are coloured according to the labels, and the nucleated actin filament subunits are coloured grey and are labelled Ac1 and Ac2. b. Overview of the SPIN90 dimer model with SPIN90 and SPIN90* coloured in magenta and pink respectively. The three α-helices (residue 312-362) from the two SPIN90s mediate the dimerisation, referred to as SPIN90 Dimerisation Domain (SDD). c. SDD inserts into a hydrophobic groove formed by the SDD from the opposite SPIN90. Hydrophobic residues on the interaction surface of one SPIN90 are labelled. Hydrophobic surface regions of the opposite SPIN90 are coloured in yellow and hydrophilic surface regions of the opposite SPIN90 are coloured in teal. The view is the same as in panel b.

In our reconstruction, Arp2 and Arp3, which are bound to ADP (*Extended Data Fig. 4a*), are both flattened and adopt a short-pitch conformation. Comparison of the Arp2/3 complexes in our reconstruction with those at branch junctions, confirmed that they are in an active conformation (*Extended Data Fig.4b*). The previous X-ray crystallography structure of SPIN90 (residues 269 to 722) was determined in complex with Arp2/3 in an inactive conformation^32^. The residues that form the SDD were present in the crystallised construct but were insufficiently ordered to unambiguously determine their structure and thus dimerisation was not visualised (*Extended Data Fig. 5*). Intriguingly, however, the overall organisation of the SPIN90-Arp2/3 complex within the crystallographic unit cell is similar to our dimeric structure with active Arp2/3, suggesting that SPIN90 had indeed formed dimers, but that the crystal context inhibited complex activation. This validates the idea that SPIN90 readily dimerises but also suggests that the SPIN90 dimer structure is most stable and therefore most readily visualised in complex with activated Arp2/3.

### The interaction between SPIN90 and Arp3 is essential for Arp2/3 activation

To investigate how SPIN90 dimerisation promotes Arp2/3 complex activation, we analysed the interactions between the SPIN90 dimer (with each monomer referred to as SPIN90 and SPIN90*) and one of the Arp2/3 complexes. SPIN90 and SPIN90* contact each Arp2/3 complex through two distinct interfaces (Fig. 3a, Movie. 3). The first, as previously defined, involves a primarily electrostatic interaction of the armadillo repeat domain of SPIN90 with the ArpC4 hinge helix (Fig. 3 a, b, Movie.3). Our structure shows that the ArpC4 hinge helix conformation matches that in the branch junction structure (*Extended Data Fig. 4b*) but is bent compared to that in the SPIN90-inactive Arp2/3 co-crystal structure (*Extended Data Fig. 5*). This shows that bending of the ArpC4 helix is a common feature of Arp2/3 complex activation, both at branches through mother filament interaction and in linear filament nucleation by SPIN90 (Fig. 3b).

**Figure 3.**
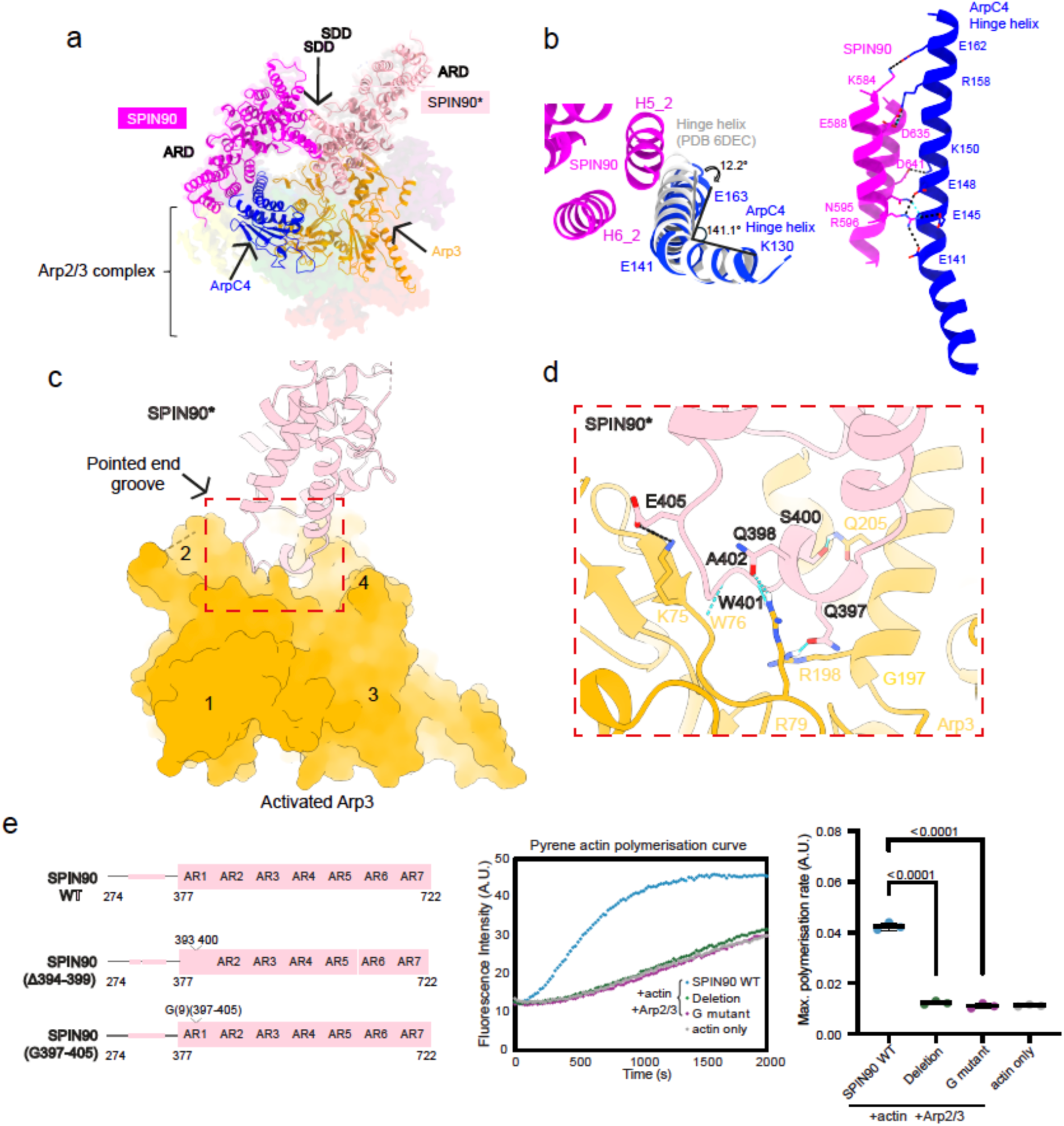
Dimeric SPIN90 forms two interfaces with each activated Arp2/3 complex to promote actin nucleation. a. Overview of two major interactions between the SPIN90 dimer and a single Arp2/3 complex. The SPIN90 dimer, ArpC4 and Arp3 are modelled and shown in ribbon representation, within the cryo-EM map. The cryo-EM map is displayed with transparency to highlight the interactions. The colour scheme is the same as in Fig. 2a. b. Comparison of the ArpC4 hinge helix in the activated state and the inactive state (PDB 6DEC). Structures are aligned based on ArpC4 residues 2-141. The ArpC4 bending angle is defined using K130 Cα, E141 Cα and E163 Cα (left). Interactions between SPIN90 (H5_2, the second helix within the fifth armadillo repeat, and H6_2, the second helix within the sixth armadillo repeat) and the ArpC4 hinge helix. Residues forming hydrogen bonds (blue dotted line) and the salt bridge (black line) are shown in stick representation (right). c. Overview of the Interaction between SPIN90* and Arp3. SPIN90 is shown in the ribbon representation while Arp3 is displayed in the surface representation. The subdomains in Arp3 are labelled. d. Zoomed-in view on the interface between SPIN90* and Arp3 pointed end groove inside the red box in panel c. Residues forming hydrogen bonds (blue dotted line) and the salt bridge (black line) are shown in stick representation. e. Schematic (left) of SPIN90 Arp3 interaction mutants, representative assay curves (middle) and quantification (right) of the maximum polymerisation rate for pyrene assay reactions in the presence of Arp2/3 complex and actin The data for actin alone corresponds to the maximum spontaneous polymerisation rate. Εach point represents the maximum polymerisation rate of a pyrene curve. The bar indicates the mean of three repeats and the error bar shows the standard deviation. Two-tailed unpaired t-test has been applied to analyse the statistical significance with p-values shown on the top.

The second, previously uncharacterised, interface forms between the N-terminal end of the armadillo repeats of SPIN90* and Arp3 (Fig. 3a, Movie.3). In this second interface, the interacting loop from SPIN90* (residues 393-406) inserts into the pointed-end groove of activated Arp3, forming electrostatic and hydrogen bonds (Fig. 3c, d). Moreover, the interaction between SPIN90* and Arp3 is unique to the activated conformation of Arp3 (*Extended Data Fig. 6a*). This is reminiscent of the role of the mother filament in stabilising the active conformation of Arp3 during branch formation (*Extended Data Fig. 6b*) which in both cases promotes binding by the first actin subunit of the nucleated filament^23^. Thus, we hypothesise that this second SPIN90 interaction stabilises Arp3 in an activated state, enabling the Arp2/3 complex to nucleate an actin filament (Movie.4).

To test this hypothesis, we generated two SPIN90 mutants: a deletion (Δ394-399) that shortens the Arp3 interaction loop and a glycine substitution (G397-405) that alters the structural properties of the interacting loop (Fig. 3e). Mass photometry confirmed that both mutants still form dimers (*Extended Data Fig. 6c*) but neither mutant could activate the Arp2/3 complex (Fig. 3e). These findings demonstrate that the interaction of the armadillo repeat domain of SPIN90 with ArpC4 and active Arp3 are essential for Arp2/3 complex activation.

### SPIN90 dimers and branch junctions bind active Arp2/3 via a similar mechanism

Comparison of the interface of activated Arp2/3 bound to the SPIN90 dimer with that of Arp2/3 on the mother filament at branch junctions reveals striking similarities in subunit contacts (Fig. 4a, white and grey surfaces). This demonstrates that the previous proposal that SPIN90 stabilises activated Arp2/3 via a very small interaction with ArpC4 is incomplete. Rather SPIN90 dimerisation generates a two-fold increase in the interaction area that stabilises activated Arp2/3. Our observations demonstrate that both branch and single filament nucleation are mechanistically more similar than had been previously appreciated in that they involve interactions with both ArpC4 and Arp3.

**Figure 4.**
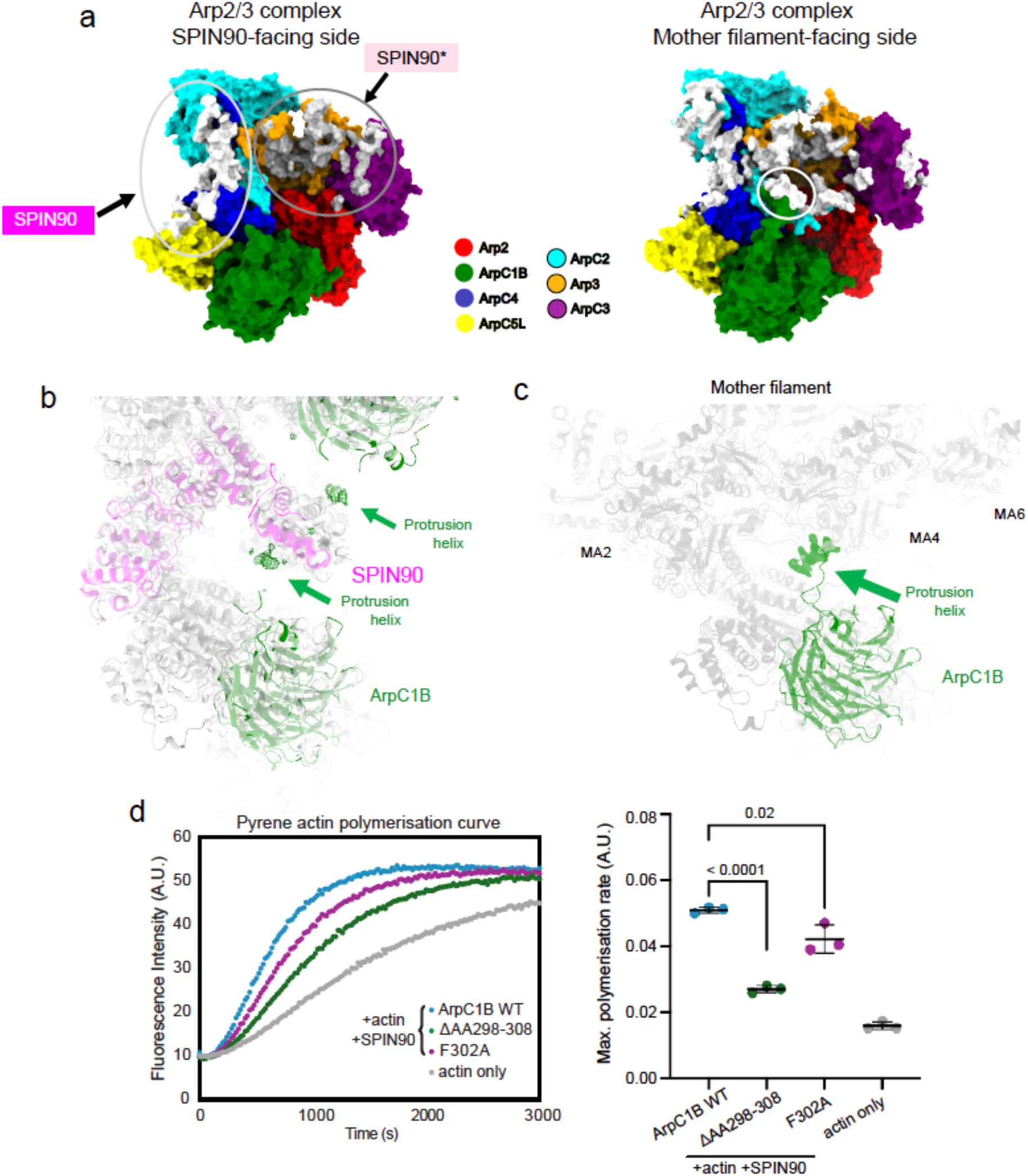
SPIN90 dimers and branch junctions bind active Arp2/3 via a similar mechanism. a. Interaction surface comparison. The interfaces are defined by residues that are within 5 Å distance of SPIN90 dimer or the mother filament (PDB 8P94). The SPIN90 interface is coloured in white and enclosed within the light grey circle, while the SPIN90* interface is coloured in grey and enclosed within the grey circle. The mother filament interface is coloured in white, with the interface on ArpC1B protrusion helix highlighted in the white circle. b. Cropped zoomed-in view of the cryo-EM reconstruction of the SPIN90-Arp2/3 complex nucleated bidirectional actin filaments. The cryo-EM map is shown with transparency. The model is fitted into the cryo-EM map. ArpC1B subunit is coloured in green, SPIN90 is coloured in magenta and other models are shown in light grey. The density attributed to the ArpC1B protrusion helix is represented as a difference in density. This difference density, shown in mesh representation, was calculated by subtracting the simulated 5 Å resolution density of the model from the cryo-EM reconstruction using the ChimeraX “volume subtract” command. c. Cropped zoomed-in view of the cryo-EM reconstruction of the cortactin-bound actin branch junction. The ArpC1B protrusion helix density is pointed with the green arrow. The mother filament subunits are labelled with MA2, MA4 and MA6. d. The representative curves (left) and quantification of the maximum actin polymerisation rate (right) for pyrene assay reactions containing the wild-type Arp2/3 complex, the Arp2/3 complex with the ArpC1B protrusion helix deleted, and the Arp2/3 complex containing the ArpC1B(FA) mutant, in the presence of SPIN90. Each point represents the results of an independent experiment. The mean of the maximum polymerisation rate and the standard error are shown. An unpaired t-test was used to calculate the p-values, which are displayed in the figure.

One point of difference between the interaction surface of Arp2/3 with the mother filament at branches compared to the SPIN90 dimer is that the ArpC1B protrusion helix, which contacts the mother filament is not apparent in the SPIN90 complex. On closer inspection, however, our structure revealed the presence of lower resolution density contacting the SDD that we attribute to the ArpC1B protrusion helix (Fig. 4b and *Extended Data Fig. 7a*). The flexibility of the long loop in which the protrusion helix is embedded would allow its contact with SPIN90 (Fig. 4b). In addition, the position and properties of the predicted ArpC1B protrusion helix would enable an interaction with a hydrophobic groove on the surface of the SDD (*Extended Data Fig. 7b*).

In the branch junction structure, the ArpC1B protrusion helix inserts a conserved phenylalanine into the hydrophobic groove of the mother filament (*Extended Data Fig. 7c*)^23^. To investigate whether the contact of the ArpC1B protrusion helix with the SDD is required for SPIN90 to activate Arp2/3, we generated complexes lacking the helix or with an F302A mutation. We found that Arp2/3 complexes lacking the ArpC1B protrusion helix had significantly reduced actin nucleating activity (Fig. 4d). The F302A mutant also showed reduced activity albeit to a lesser extent, consistent with the idea that other residues in the protrusion helix contribute to its interaction with the SDD (Fig. 4d). Taken together, our observations suggest that the ArpC1B protrusion helix also functions in SPIN90-mediated activation of Arp2/3 to increase the interaction between SPIN90 and the Arp2/3 complex, albeit in a more flexible and dynamic way compared with the mother filament.

### Species-specific differences in Arp2/3 activated by the WISH/DIP/SPIN90 family

Having investigated the mechanism by which the SPIN90 dimer stabilises activated mammalian Arp2/3, we compared our SPIN90-bound Arp2/3 structure with monomeric yeast Dip1-Arp2/3 and with reference to Arp2/3 at actin branches. A striking difference between the SPIN90-activated human Arp2/3 and the Dip1-activated yeast complex relates to the flattening of Arp3^22, 23, 36^. In our SPIN90-activated human complex, both Arp2 and Arp3 are fully flattened with a dihedral angle of −2.5° for Arp3, that represents a fully active state primed for nucleation (*Extended Data Fig. 8a*, Fig. 5a). In contrast, the Dip1-activated yeast Arp2/3 exhibits partial flattening of Arp3, with a dihedral angle of −9.3° ^36^. The complete flattening seen in our structure can be explained by the additional interaction between the SPIN90 dimer and Arp3. In addition, in the Dip1-actived Arp2/3 structure, ArpC3 does not contact Arp2 (*Extended Data Fig. 8b*, purple arrow), whereas in the SPIN90-activated complex, ArpC3 connects Arp2 and Arp3, likely due to the full flattening of Arp3. Moreover, the extended N-terminus of human ArpC5L inserts between Arp2 and Arp3 in the SPIN90-activated Arp2/3 structure in contrast to the shorter yeast ArpC5 N-terminus that only contacts Arp2; these differences are also observed in the actin branch junction structures (*Extended Data Fig. 8b*, yellow arrow)^22, 26^. Overall, our analyses indicate general differences in the properties of Arp2/3 complexes from human and yeast that are reflected in mechanisms by which they are activated.

**Figure 5.**
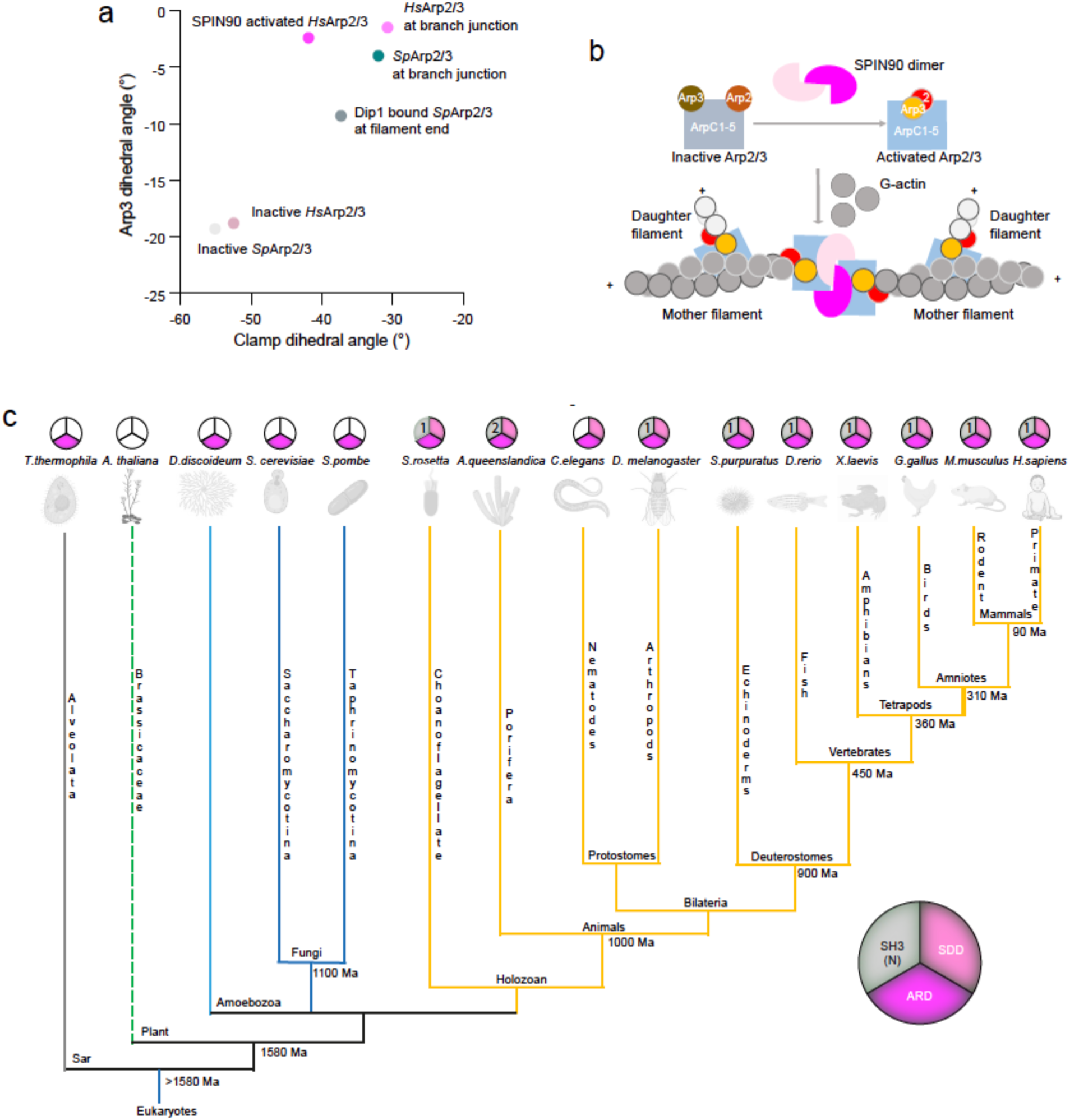
Conservation of dimerisation in metazoan SPIN90 orthologues suggests a common mode of anti-parallel actin filament formation. a. Right: Plot of the clamp dihedral angle vs Arp3 dihedral angle in representative published Arp2/3 structures (*Extended Data Fig. 8a*). The clamp-twist angle was defined as the dihedral between K18 Cα (ArpC2), I244 Cα (ArpC2), S147 Cα (ArpC4) and R32 Cα (ArpC4) in the *Hs.* Arp2/3 complex or between R18 Cα (ArpC2), I262 Cα (ArpC2), S147 Cα (ArpC4) and R32 Cα (ArpC4) in the *Sp.* Arp2/3 complex. Arp3 dihedral angle is identified as the dihedral between the centres of mass of subdomain 2, subdomain 1, subdomain 3 and subdomain 4. b. Schematic showing bidirectional mother filament organisation enables more spread out daughter filament organisation. c. Phylogenetic analysis of SPIN90 homologues in 15 eukaryotes model organisms. The pie chart illustrates the domain organisation of human SPIN90, divided into 3 sections representing the three major domains (SH3 domain, SPIN90 Dimerisation Domain (SDD), armadillo repeat domain (ARD)) identified using AlphaFold predictions. A coloured slice indicates the presence of the corresponding domain, while an uncoloured slice indicates its absence. The number (N) in the SH3 domain section specifies the number of SH3 domains present. The phylogenetic tree indicates the model organisms’ relationships and estimated divergence time (Ma, million years ago)^61^. Please note that the branch lengths do not reflect evolutionary time.

## DISCUSSION

Using *in vitro* reconstitution and biophysical analyses, we discovered that human SPIN90 forms a dimer that activates two Arp2/3 complexes to nucleate bidirectional actin filaments. Moreover, our analysis demonstrates that the mechanisms by which mammalian Arp2/3 binds mother filaments at branches and at the end of linear filaments are more similar than have been previously thought. The SPIN90 dimer forms two interfaces with each Arp2/3 complex, binding to both the bent ArpC4 hinge helix and the pointed end of activated Arp3. We infer that dimerisation, by increasing the total buried surface in the complex, contributes to stabilisation of the activated state of human Arp2/3 complex. In contrast in the yeast Arp2/3 complex, which has a lower energy barrier for activation^18, 19, 38^, we suggest that both interactions are not required. The higher energy barrier for human Arp2/3 activation also explains why SPIN90 has evolved to form a dimer to enhance its activation ability compared to its yeast counterpart Dip1. These observations emphasise the structural plasticity of the Arp2/3 complex between human and yeast and show how WISH/DIP/SPIN90 family proteins - and presumably other Arp2/3 regulators - have evolved to accommodate the conformational variability of their cognate Arp2/3 complexes.

If the bidirectional actin filaments we observed were to act as mother filaments for subsequent actin branch initiation and propagation, then their bidirectionality would have significant implications for the architecture of the resulting actin network. First, the organisation of the SPIN90 dimer is predicted to position the SH3 and poly-proline domains of each protein on the same side of the nucleating complex (*Extended Data Fig. 2b*). Such an arrangement would facilitate one-sided interactions with signalling cascades at the plasma membrane, potentially aiding the activation process. Second, the multivalent interactions between Nck, NPFs, and SPIN90 would give the branch formation process a robust jump start^30^. Once the mother filaments are nucleated, the Class 1 NPFs can immediately promote the daughter filament assembly. Interestingly, Nck could interact with both SPIN90 and N-WASP via its three SH3 domains and thereby couple linear and branched actin filament nucleation^39, 40^. Lastly, bidirectional organisation can enable efficient actin branch formation as daughter filaments can be nucleated in opposite orientations, distributing the force for cellular processes such as cell migration and phagocytosis (Fig. 5b) ^41^. The overall flexibility of SPIN90-Arp2/3 complex (Movie 2) may also contribute to the stability of SPIN90-Arp2/3 nucleated filaments in the presence of piconewton forces^42^. Future *in vitro* and cell-based studies together with mathematical modelling will be required to determine precisely how SPIN90-mediated bidirectional mother filament organisation might enhance the efficiency of branched actin propagation, membrane modulation and response to external forces.

The differences in the structures of Dip1 and SPIN90-activated Arp2/3 complexes between human and yeast and their activation mechanisms prompted us to investigate conservation of the SPIN90 family more broadly. Comparison of SPIN90 homologues across 15 eukaryotic model organisms allowed us to identify distinct patterns that highlight the evolutionary trajectory of this protein family (Fig. 5c). Notably, no SPIN90 family proteins were identified in *Arabidopsis*, indicating a potential evolutionary loss or replacement by alternative functional analogues in plants. Intriguingly, however, SPIN90 orthologues in all examined metazoans contain an SDD, together with SH3 and armadillo repeat domains in a conserved N- to C-terminal order; the exception is *C. elegans*, which lacks an SH3 domain. Furthermore, AlphaFold structural predictions of the SDD and armadillo repeat domains of these proteins indicate these metazoan SPIN90 orthologues are likely to form dimers (*Extended Data Fig. 9*). Interestingly, even SPIN90 from the unicellular holozoan protist *S. rosetta,* a choanoflagellate and the sister group of metazoans, also contains all three domains^43^. In contrast, two fungal species and other protists, such as *T. thermophila* and *D. discoideum* (slime mould) are predicted to contain armadillo repeat domains, within which the previously identified leucine-rich domain is located, but the SDD and SH3 domain are absent. This structural divergence reflects functional specifications aligned with the complexity of signalling networks in different evolutionary lineages. Indeed, the conservation of the three-domain and dimeric arrangement in choanoflagellates suggests that it is an ancient evolutionary adaptation specific to the holozoan clade, occurring before the emergence of multicellular animals^44^. This hints at an ancient evolutionary path that laid the foundation for the complex actin system observed in animals today. In the future, it will be important to explore the implications of SPIN90 dimerisation in its regulation and cellular role in specifying bidirectional actin filament polymerisation. This has the potential to affect the dynamics and geometry of the resulting actin networks in different cellular contexts.

## Acknowledgements

This project has received funding from the European Research Council (ERC) under the European Union’s Horizon 2020 research and innovation programme (grant agreement No 810207 to M.W. and C.A.M.). L.C. was supported by the European Union’s Horizon 2020 Marie Sklodowka-Curie individual fellowship program (H2020-MSCA-IF-101028239 – MolecularArp). G. R.-L. was supported by the French Agence Nationale de la Recherche (grant ANR-21-CE13-0043). M.W. is supported by the Francis Crick Institute, which receives its core funding from Cancer Research UK (CC2096), the UK Medical Research Council (CC2096), and the Wellcome Trust (CC2096). We acknowledge Diamond Light Source for access and support of the cryo-EM facilities at the UK national Electron Bio-imaging Centre (eBIC) proposal Bl34130. We thank Natalya Lukoyanova and Shu Chen for electron microscope support and David Houldershaw for computing support at Birkbeck. We thank Richard Mitter for advice about the phylogenetic analysis, Roger George for mass photometry support and Antoine Jégou (Institut Jacques Monod) and Giulia Zanetti (Birkbeck, University of London) for feedback on the manuscript.

## Author contribution statement

T.L purified recombinant capping protein, conducted the cryo-EM sample preparation, data collection and structural analysis; L.C. purified all other recombinant Arp2/3 regulators and designed, conducted and analysed in vitro biophysical experiments and the TIRF-M assay; M.M. prepared recombinant Arp2/3 complexes; G R-L provided access to and technical advice on TIRF-M experiments; M.W. and C.A.M. supervised the project; T.L., M.W. and C.A.M. wrote the paper, with input from all authors.

## METHODS

### Protein purification

Human SPIN90-C (residues 274 - 722) and its mutants were purified following the protocol described by Cao et al^42^. Human Arp2/3 complex (containing ArpC1B and ArpC5L) was purified following the protocol by Baldauf et al^45^. Mouse capping protein α1β2 was purified following the protocol described by Liu et al and some proteins used in that previous study were reused here^26^.

### Mass photometry

Purified recombinant SPIN90-C or its mutants was diluted to 30 nM with PBS (Phosphate Buffered Saline). Large aggregates were eliminated by centrifuging at 21,130 rcf for 10 min at 4°C. 2 µL of diluted protein was then added to 18 µL of PBS in the well of a gasket on a TwoMP instrument (Refeyn) and events were recorded with AquireMP software (Refeyn) at room temperature. BSA (66 kDa, Thermo Scientific™, #23209), ADH (150 kDa, Sigma-Aldrich, #A7011) and Urease (90/272/544 kDa, Sigma-Aldrich, #94280) were used as standards.

### TIRF microscopy assays

Deep UV-treated coverslips were passivated with mPEG silane overnight and thoroughly rinsed with ethanol and water. Flow chambers were prepared as described in Cao et al^46^. G-actin (0.5 µM, 15% labelled with Alexa-488) was pre-incubated with SPIN90 and the Arp2/3 complex for 4 minutes in the incubation buffer containing 5 mM Tris-HCl (pH 7.0), 50 mM KCl, 1 mM MgCl_2_, 0.2 mM EGTA, 0.2 mM ATP, 10 mM DTT, and 1 mM DABCO at room temperature. Alternatively, 1 µM G-actin (15% labelled with Alexa-488) was incubated for over 10 minutes in the incubation buffer to generate spontaneously nucleated actin filaments.

Pre-polymerised actin was then mixed with 0.5 µM G-actin (15% labelled with Alexa-568) in the imaging buffer, which included 0.1% BSA and 0.3% methylcellulose in addition to the incubation buffer and loaded directly onto the TIRF microscope. Image acquisition was performed at 25°C. For each independent repeat, the fraction of filaments with double barbed ends was quantified. The mean and standard deviation were then calculated and plotted.

Fiji software^47^ was used to analyse images manually. To randomly select actin filaments for analysis, the red channel was turned off to avoid biasing filament selection according to their growth state. Pre-polymerised filaments with strong 488 nm signals were excluded, as they were most likely actin bundles with uncontrollable numbers of filament ends. Prism 10 (10.1.1) software (GraphPad) was used to calculate all the statistical analysis.

### Cryo-EM sample preparation

Porcine βλ non-muscle actin lyophilised powder (Hypermol, #8105-01) was dissolved in water to obtain a stock solution of 1mg/ml (23.8 μM). To reconstitute SPIN90-C, the Arp2/3 complex, and their nucleated actin filaments, reconstitution conditions were adapted from that used to generate actin branches with the exclusion of VCA and cortactin^26^. Specifically, 1.5 μM Arp2/3 complex, 14.3 μM SPIN90-C, 0.7 μM actin and 2.9 μM capping protein were mixed in 16.8 μL reaction buffer (20 mM HEPES pH 7.5, 50 mM KCl, 1 mM EGTA, 1 mM MgCl2, 0.2 mM ATP and 1 mM DTT). The mixture was incubated at room temperature for 20 min. Then 0.5 μL of 23.8 μM actin stock was added in 9 steps. After each addition of actin, the mixture was incubated at room temperature for 20 min. Additionally, 0.6 μL of 80 μM capping protein was added together with the third and seventh addition of actin. At the end of the reaction, 100 μM phalloidin was added to stabilise the polymerised actin filaments.

4ul of the mixture was applied to a glow-discharged C flat 1.2/1.3 Cu grid. The grid was plunge frozen into liquid ethane using EM GP2 Automatic Plunge Freezer (Leica). The sensor and back blotting parameters are as follows: additional movement of 0.3mm, blotting time of 5s, temperature of 25 °C and humidity of 95%.

### Cryo-EM data acquisition

Cryo-EM data was collected on a Titan KriosIV (Thermo Fisher Scientific) at the Diamond electron Bio-Imaging Centre (eBIC) equipped with a K3 detector and a BioQuantum energy filter (Gatan). The microscope was operated at an acceleration voltage of 300 kV with a nominal magnification of 81,000x and a pixel size of 1.06 Å. 12,512 movies were collected using EPU with the following parameters: super-resolution mode, a dose rate of 22.0 e/pixel/s, an exposure time of 2s, 50 frames and a defocus range of −0.9 to −2.4 μm.

### Cryo-EM data processing

Cryo-EM data was processed using CryoSPARC^48^. Movies were motion-corrected using Patch motion correction. CTF parameters were determined using Patch CTF. A total of 10,235 micrographs with a CTF fit resolution better than 8 Å and total frame motion distance less than 45 pixels were selected for further data processing.

Initially, 2,138,010 particles were selected using Blob picker, with a minimal particle diameter of 150 Å and maximum particle diameter of 200 Å. These particles were extracted with a binning factor of 4. Multiple rounds of 2D classification were used to remove junk particles (e.g. carbon, ice) and actin filament segments. The remaining 87,216 particles were subjected to ab-initio reconstruction with three classes. 13,727 unbinned particles from class 3 displaying the characteristic ArpC1B β propeller density were selected for homogeneous refinement, obtaining a reconstruction at 4.2 Å.

To further improve the resolution, the particles used in the initial reconstruction were subjected to another round of 2D classification to remove residual junk. The refined particle set was used for Topaz training. Using the obtained Topaz model, 26,606 and 39,254 particles were picked from the 1st and 2nd halves of the dataset, respectively^49^. These two particle sets were subjected to ab initio reconstruction.

Particles from the class resembling the previous homogeneous refinement reconstruction were combined and subjected to multiple rounds of 2D classification to remove junks. Finally, the remaining 39,104 particles were refined using NU-refinement, and the resolution was further improved after applying C2 symmetry. Global and local resolutions were estimated in CryoSPARC.

Particles from the final C1 reconstruction were used for 3D variability analysis in CryoSPARC with six modes to solve and a filter resolution of 8 Å^50^. 3D Variability result was displayed in cluster mode with 4 clusters, a var_range_percentile of 0 % and a filtered resolution of 6 Å. Two reconstructions from the particle clusters (cluster 1 and 4) with the most divergent component values were used for model building.

### Model building

The Arp2/3 complex and four daughter filament subunits from the PDB 8P94, along with the SPIN90 structure from PDB 6DEE, were rigidly fitted into one asymmetric unit of the cryo-EM map using ChimeraX, followed molecular dynamic-based flexible fitting using ISOLDE^51, 52^. Namdinator was used to optimise bond geometry^53^. Clashes, Ramachandran outliers and rotamer outliers were manually corrected using ISOLDE and Coot^54^. Phenix real space refinement was used for refining the model^55^. Any residual clashes, Ramachandran outliers and rotamer outliers were again manually fixed using ISOLDE and Coot. The model was then duplicated and fitted into the density of the other asymmetric unit. Models used in 3D variability analysis in Movie 2 were further refined from the above model using Namdinator and morphed using ChimeraX.

### Phylogenetic analysis

Orthologues of SPIN90 in 15 model organisms were identified using InterPro (IPR30125) and Ensembl^56, 57^. Using predictions from the AlphaFold database, the presence of SH3 domain, SDD and armadillo repeat domains in these proteins was analysed^58, 59^. The dimeric structures of the representative SPIN90 family protein were predicted using the AlphaFold server^60^.

### Pyrene assay

To test the nucleation efficiency of Arp2/3 ± SPIN90, 8 nM Arp2/3 C1B/C5L complex, 400 nM SPIN90, and 2.5 µM G-actin (5% pyrene-labelled) were mixed and measured in a Safas Xenius fluorimeter at room temperature. The negative control was done by simply mixing the same amount of pyrene actin and SPIN90 without Arp2/3 complex. The experimental buffer contained 5 mM Tris-HCl (pH 7.0), 50 mM KCl, 1 mM MgCl2, 0.2 mM EGTA, 0.2 mM ATP, 10 mM DTT, and 1 mM DABCO, all maintained at room temperature.

For each experimental condition, the pyrene assay was repeated three times independently. The maximum polymerisation rate for each pyrene curve was measured and plotted. The mean and standard deviation for each condition were calculated. A two-tailed unpaired t-test was applied to analyse the statistical significance, with p-values shown in the figures.

## DATA AVAILABILITY

The cryo-EM reconstruction is deposited in the Electron Microscopy Data Bank under the following accession codes: EMDB-xxxxx. The corresponding structural model is deposited in the Worldwide Protein Data Bank under the accession code PDB ID: XXXX.

**Movie 1 - Cryo-EM reconstruction of SPIN90-Arp2/3 complex nucleated actin filaments**

**Movie 2 - 3D variability analysis reveals the structural elasticity of SPIN90-Arp2/3 complex**

**Movie 3 - SPIN90 dimer interacts with a single Arp2/3 complex with two major binding sites**

**Movie 4 - SPIN90 binding promotes the conformational changes of Arp3 towards activation**

## EXTENDED DATA

**Extended Data Fig. 1.**
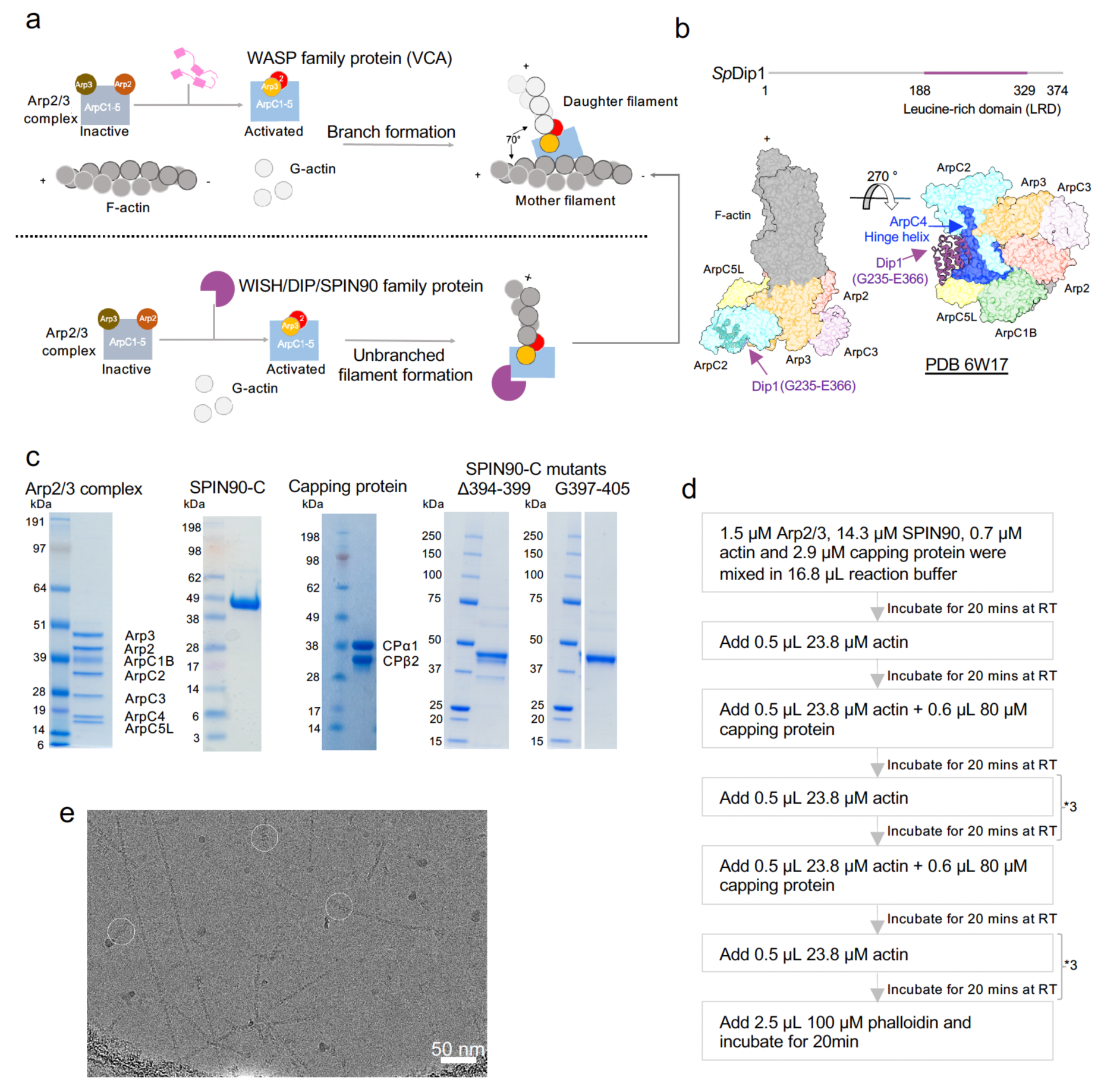
Overview of actin filament nucleation, purified proteins used in actin filament reconstitution and exemplar cryo-EM micrograph. a. Schematic of Arp2/3-mediated actin branch formation and actin branch initiation. b. Top: *Sp.* Dip1 domain organisation. Bottom: Complex structure of Dip1 and Arp2/3 complex at the end of their nucleated actin filament (PDB 6W17). The Arp2/3 complex is shown in transparent surface representation, while the resolved segment of Dip1 is shown in grape colour using ribbon representation. c. SDS-PAGE gels showing purified proteins used in cryo-EM, pyrene assay and TIRF experiments, including purified proteins that had been used in a previous study of ours^26^. d. Flow chart showing how the cryo-EM sample was prepared. e. A representative cryo-EM image of SPIN90-Arp2/3-nucleated actin filaments. Particles used for data processing are highlighted in white circles. Scale bar = 50 nm.

**Extended Data Fig. 2.**
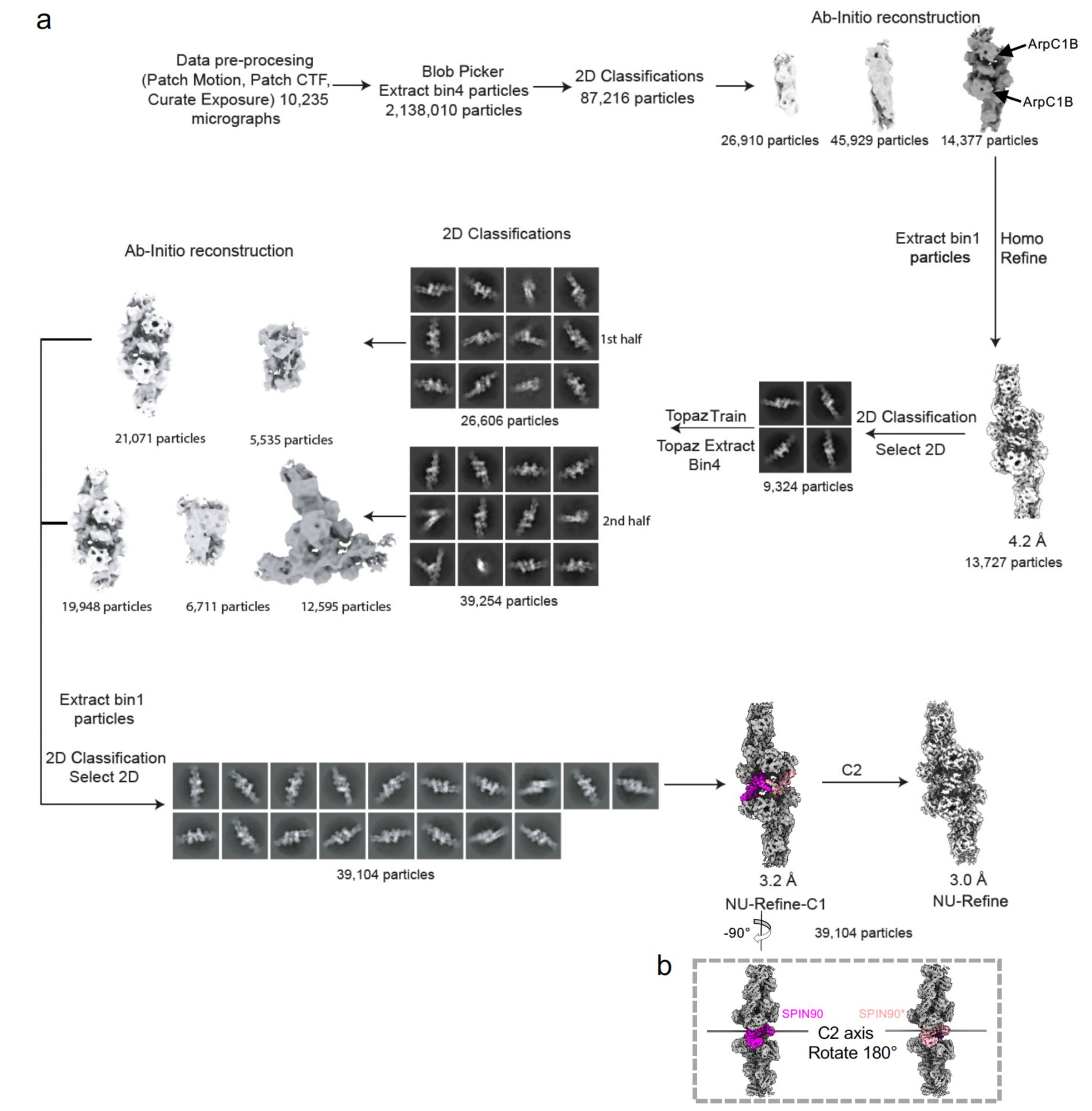
Image processing workflow for cryo-EM reconstruction. a. Data were processed using CryoSPARC. After initial pre-processing steps (Patch motion, Patch CTF and Curate exposure), 10,235 micrographs were selected for further processing. Multiple rounds of 2D classification were used to clean the dataset, followed by ab initio reconstruction with three classes. Particles from the class showing the characteristic ArpC1B β-propeller density were selected for 3D refinement. Subsequently, another round of 2D classification was used to further clean the data. The remaining 9,234 particles were used as a positive training set for Topaz particle selection. The Topaz-selected 65,860 particles were subjected to 2D classifications and Ab-initio reconstructions in two batches. A total of 39,104 particles were selected for the final reconstruction with C2 symmetry imposed. b. Two views of the C1 reconstruction with SPIN90 (magenta) and SPIN90* (pink) coloured. The C2 axis is indicated. Such an arrangement would facilitate one-sided interactions with signalling cascades at the plasma membrane via the SH3 domain and Poly-P domain, potentially aiding the activation process.

**Extended Data Fig. 3.**
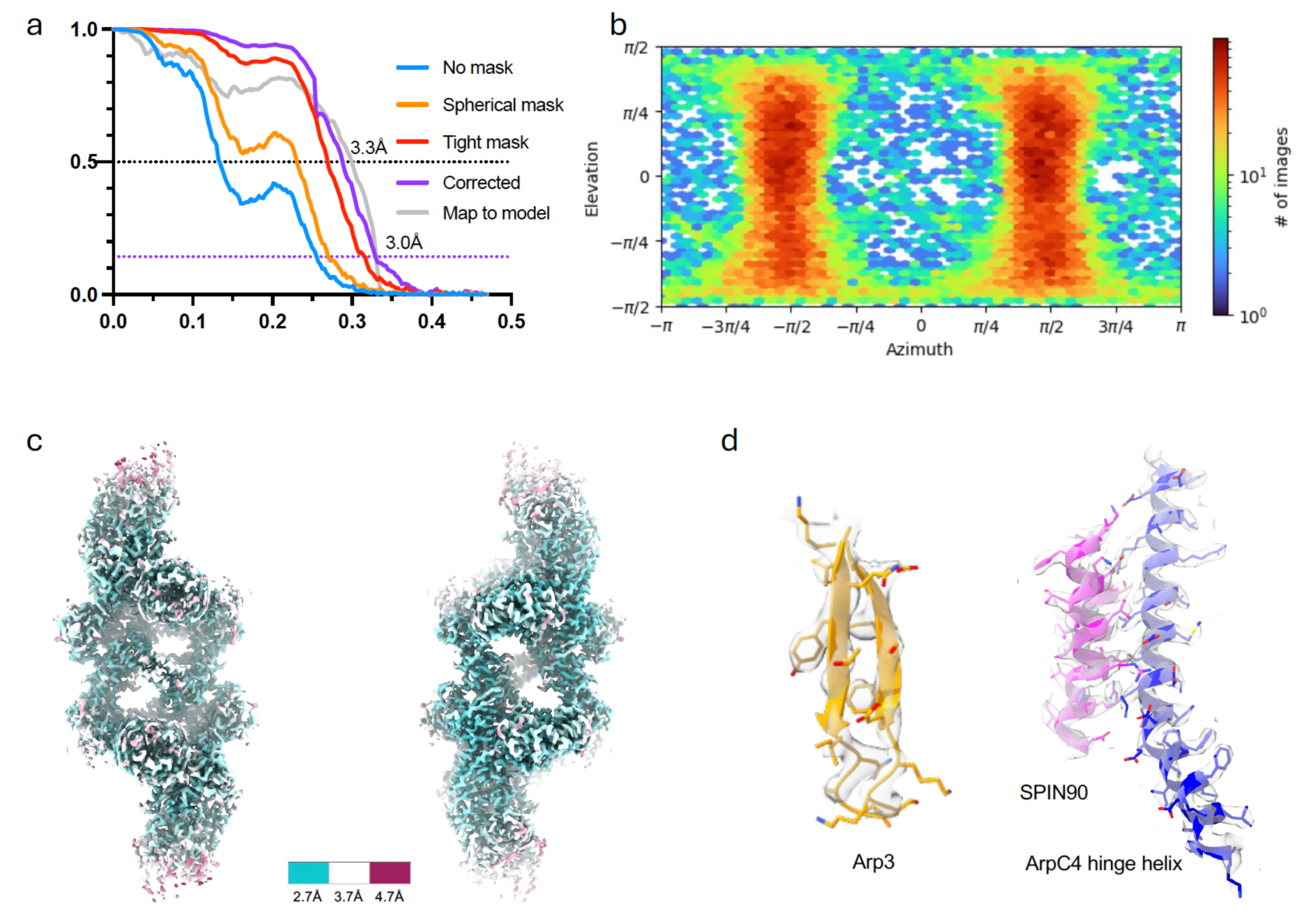
Cryo-EM data quality and validation of locally refined reconstructions. a. Half-map and map to model Fourier Shell Correlation (FSC). Global resolution (FSC cut off 0.143) and map-model resolution (FSC cut off 0.5) are indicated. b. The angular distribution of particles used for the final 3D refinement. c. Cryo-EM reconstruction coloured by estimated local resolution. d. Representative density with the fitted model showing the assignment of amino acid side chains.

**Extended Data Fig. 4.**
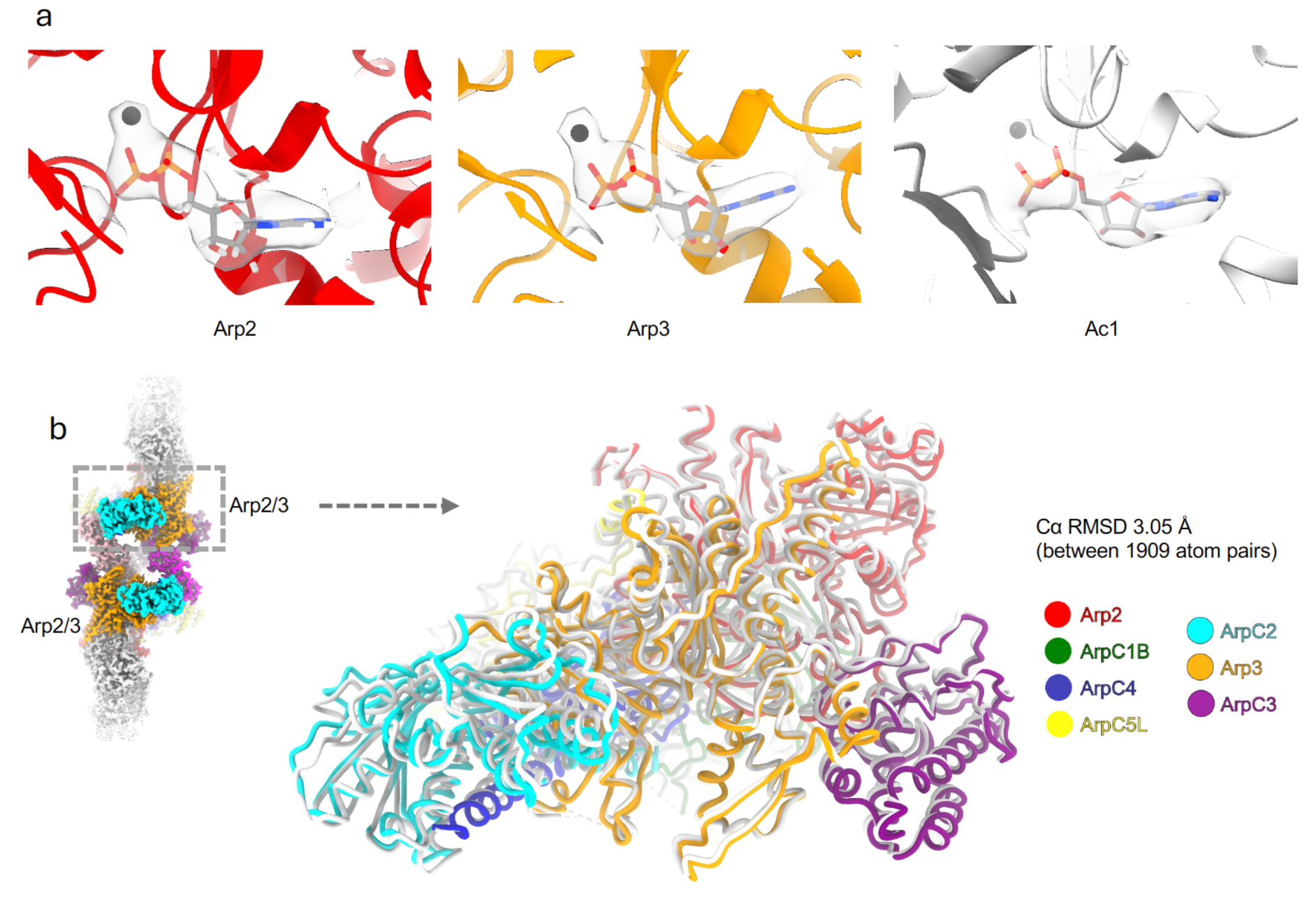
Nucleotide occupancy and demonstration of active state of the Arp2/3 complex. a. Density (transparent) from the symmetrised reconstruction and models of ADPs (in stick representation) and Mg2+ (black dot) in Arp2, Arp3 and actin subunit Ac1. b. Structural comparison of the SPIN90-activated Arp2/3 complex (this manuscript, subunits coloured) and the Arp2/3 complex at the branch junction (PDB 8P94, subunits in light grey). Structures were aligned on Arp3 subdomain 1 and 2 (AA6-37, AA60-153 and AA375-409). Overall Cα RMSD is shown.

**Extended Data Fig. 5.**
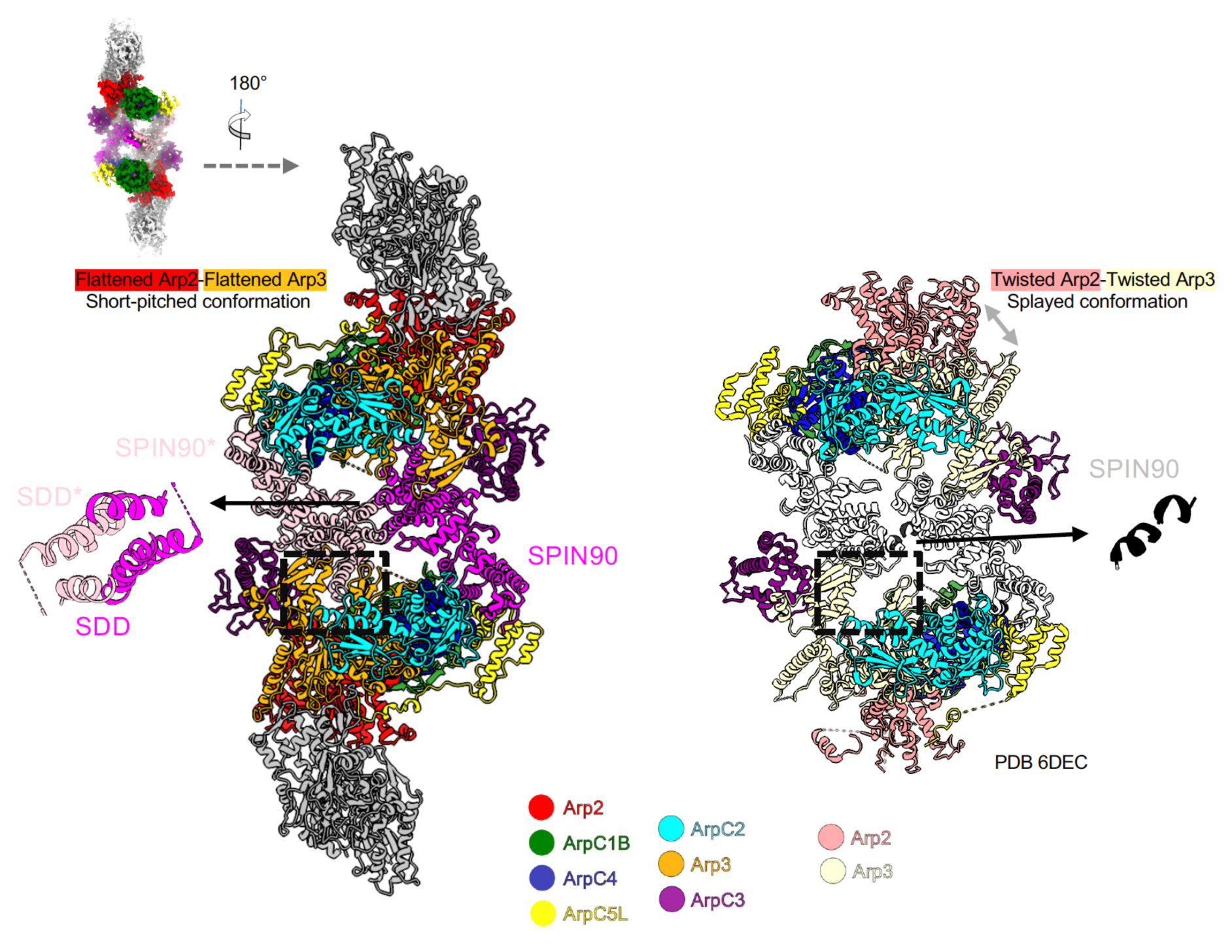
Structural comparison of the SPIN90-activated Arp2/3 complex with its nucleated actin filaments and SPIN90-inactive Arp2/3 co-crystal structure. Model of the SPIN90-activated Arp2/3 complex with its nucleated actin filaments (left). Arp2/3 subunits and SPIN90 dimer are coloured-coded as in Fig. 2a with the nucleated actin filaments shown in grey. A zoomed-in view highlights the three α-helices (SDD) from both SPIN90. Arp2 and Arp3 are flattened (activated) and arranged in a short-pitched conformation. The structural details in the black box are analysed in Fig. 3. SPIN90- inactive Arp2/3 co-crystal structure (right, PDB 6DEC). Arp2/3 subunits are coloured as in Fig. 2a, except for Arp3, which is coloured in dark salmon, and Arp2, coloured in pale goldenrod. SPIN90 is coloured in light grey. There are two short α-helices at the location of the three α-helices in the left panel, coloured in black. Arp2 and Arp3 are twisted (inactive) and adopt a spayed conformation. The structural details in the black box are analysed further in Extended Data Fig. 6a.

**Extended Data Fig. 6.**
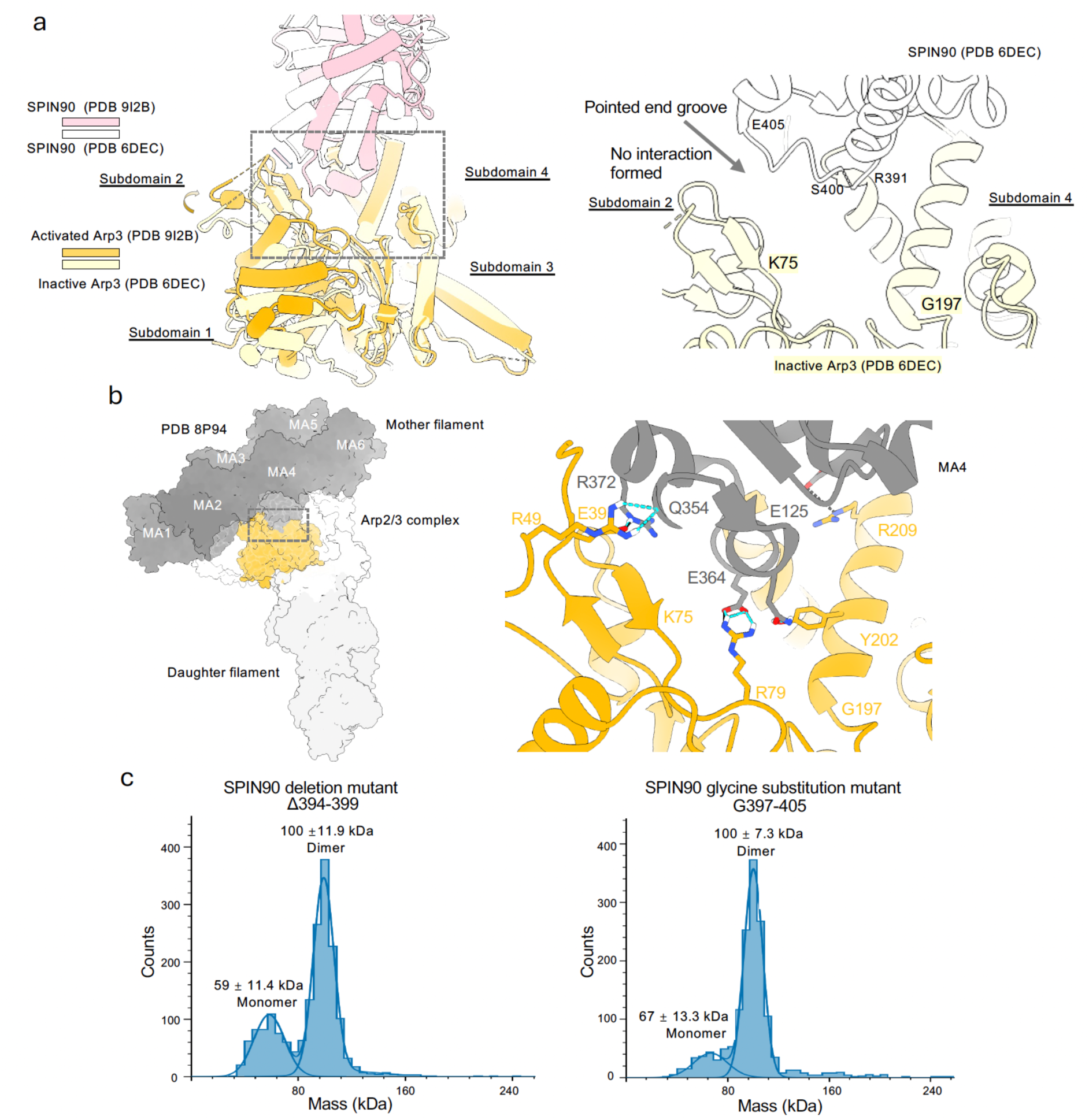
The interaction between the SPIN90 and Arp3 is unique to the activated conformation of Arp3. a. Left: Overlay of the models containing SPIN90 and Arp3 (Extended Data Fig. 5). The models are aligned on the subdomains 3 and 4 of the Arp3. Subdomains in Arp3 are labelled. Right: In the SPIN90-inactive Arp2/3 co-crystal structure (grey box on the left and black box in Extended Data Fig. 5, right panel), approximately half of the SPIN90 loop is unresolved due to its flexible nature, while the other half is positioned away from the Arp3 pointed-end groove, resulting in a loss of stable interactions. The Arp3 pointed end groove is indicated with a black arrow. End residues and breaking points of the SPIN90 loop are labelled. b. Left: Actin branch junction structure is shown in surface representation (PDB 8P94). The mother filament is coloured in dark grey. The daughter filament is coloured light grey. Arp3 is shown in orange while other Arp2/3 subunits are shown in transparency. The mother filament subunits are labelled with MA1-MA6. Right: The cropped zoomed-in view of the interaction between MA4 and Arp3. Residues forming hydrogen bonds (blue dotted line) and the salt bridge (black line) are shown in stick representation. c. Mass distribution of 30 nM SPIN90 mutant molecule, whose molecular weight are around 50 kDa respectively in theory. For the deletion mutant, two peaks were observed with molecular weight corresponding to 100 ± 11.9 kDa and 59 ± 11.4 kDa. For the G loop mutant, two peaks were observed with molecular weight corresponding to 100 ± 7.3 kDa and 67 ± 13.3 kDa.

**Extended Data Fig. 7.**
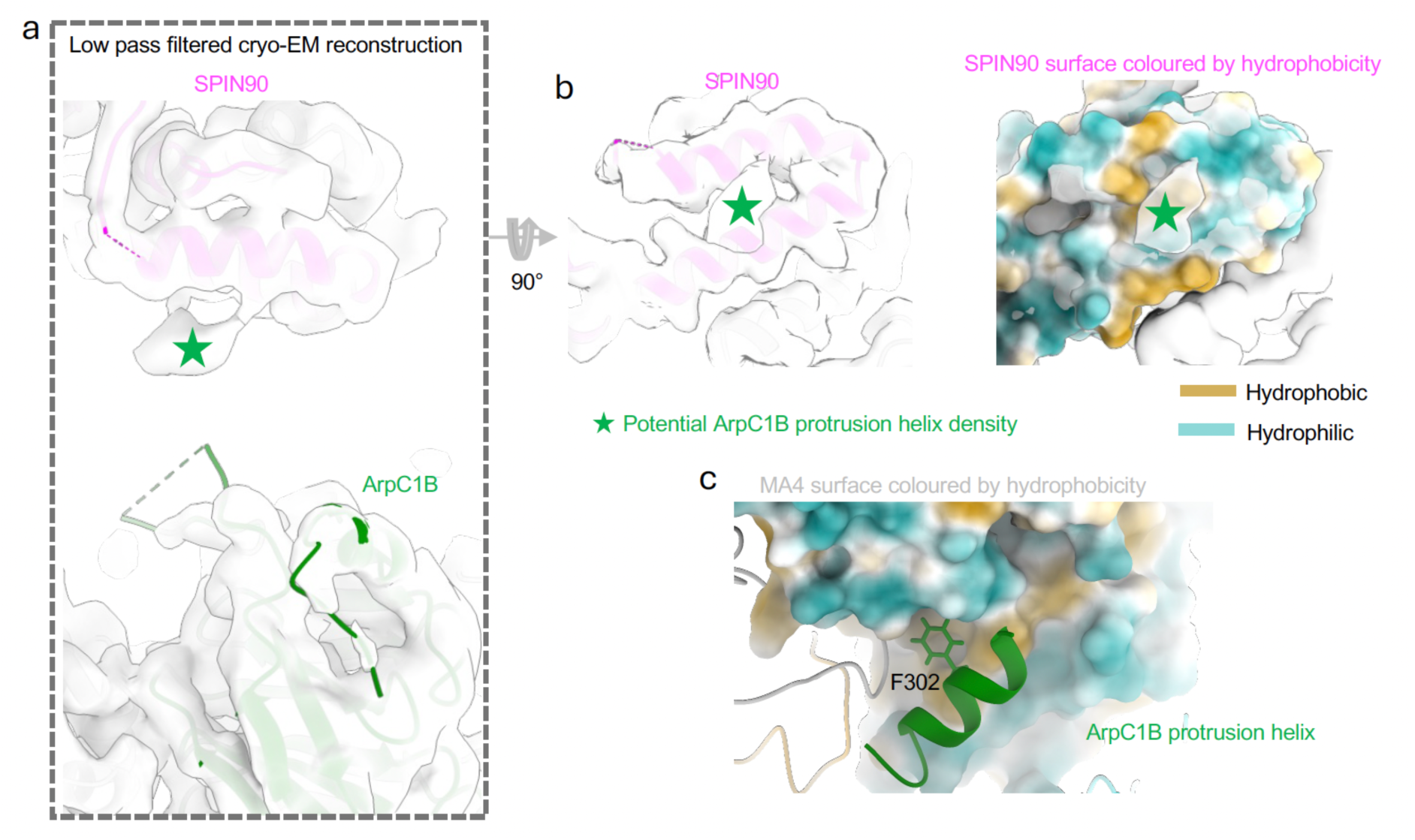
SPIN90 forms a loose interaction with the ArpC1B protrusion helix in the activated Arp2/3 complex. a. Zoomed-in view of the undefined density beneath the SDD and adjacent to ArpC1B. Reconstruction was filtered using “volume gaussian” in ChimeraX (with a standard deviation of 1Å) to reduce noise and bring out the shape of the undefined density. b. The modelled SPIN90 surface is coloured based on hydrophobicity showing a hydrophobic patch facing the potential ArpC1B protrusion helix density, highlighted with a green star. Hydrophobic regions are coloured in yellow and hydrophilic surface regions are coloured in teal. c. Surface representation of the MA4 subunit in the mother filament at the actin branch junction, coloured by hydrophobicity. The interacting ArpC1B protrusion helix is shown in the ribbon representation. The hydrophobic F302 is also highlighted with stick representation.

**Extended Data Fig. 8.**
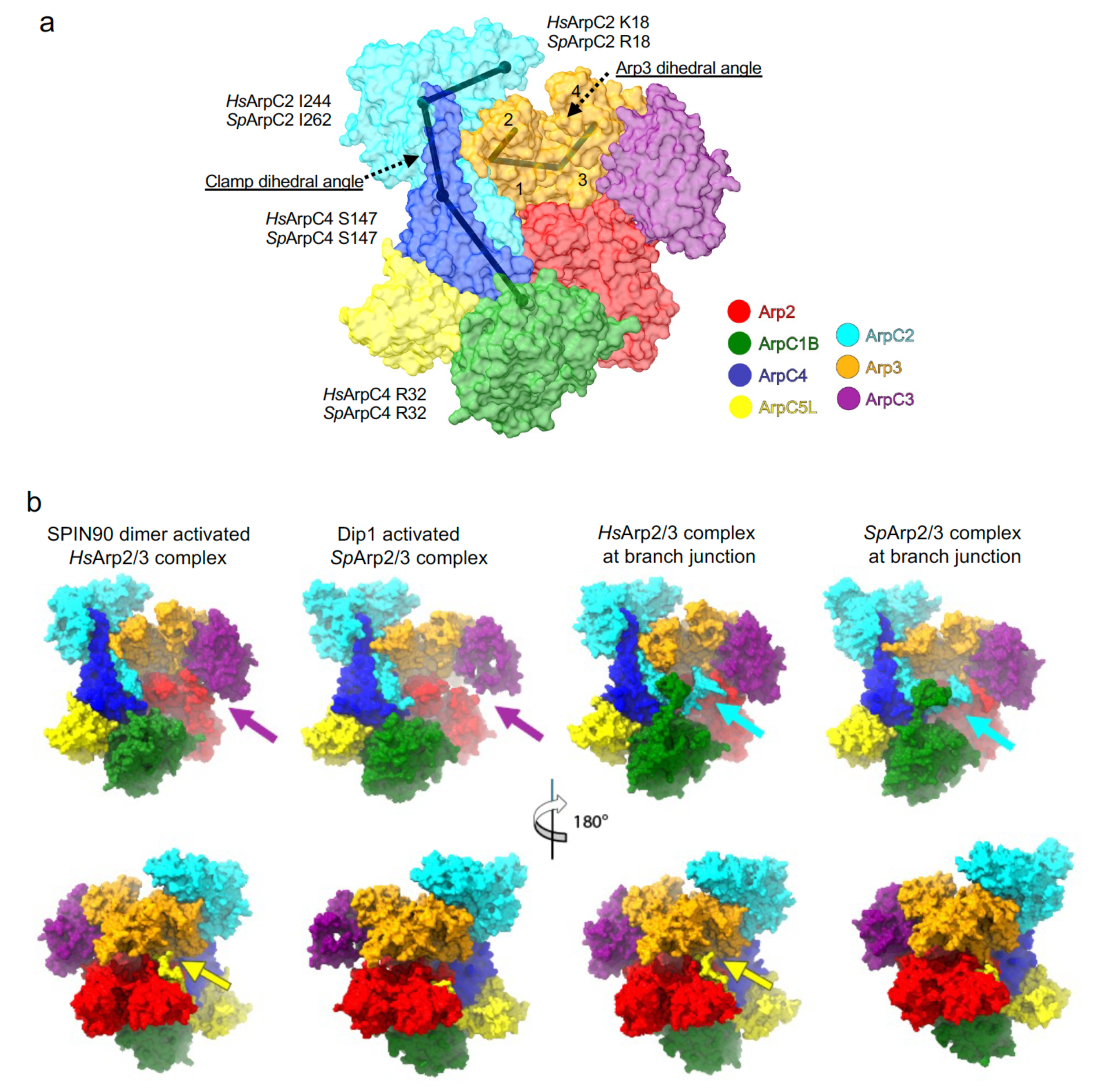
Comparison of the structures and interactions in activated human and yeast Arp2/3 complexes. a. Arp2/3 surface representation highlighting dihedral angle measurements for clamp twisting (short pitch formation) and Arp3 flattening in Fig. 5a and Extended Data Fig. 5. b. Comparison of diÖerently activated human (Hs) and yeast (Sp) Arp2/3 complexes. Arp2/3 complexes are shown in surface representation. Individual proteins are coloured according to the labels. The purple arrows highlight that the contacts between Arp2 and ArpC3 are absent in the Dip1 nucleated yeast Arp2/3 complex structure. The yellow arrows point to the insertion of the ArpC5(L) N-terminus into the groove formed by Arp2 and Arp3 only observed in human Arp2/3 complex structures. The N-terminus of *Sp*ArpC5 is 7 residues shorter and therefore cannot form these contacts. The blue arrows reveal the specific feature of ArpC2 only observed in branch junction structures, mediating the interaction of the Arp2/3 complex with the mother filament.

**Extended Data Fig. 9.**
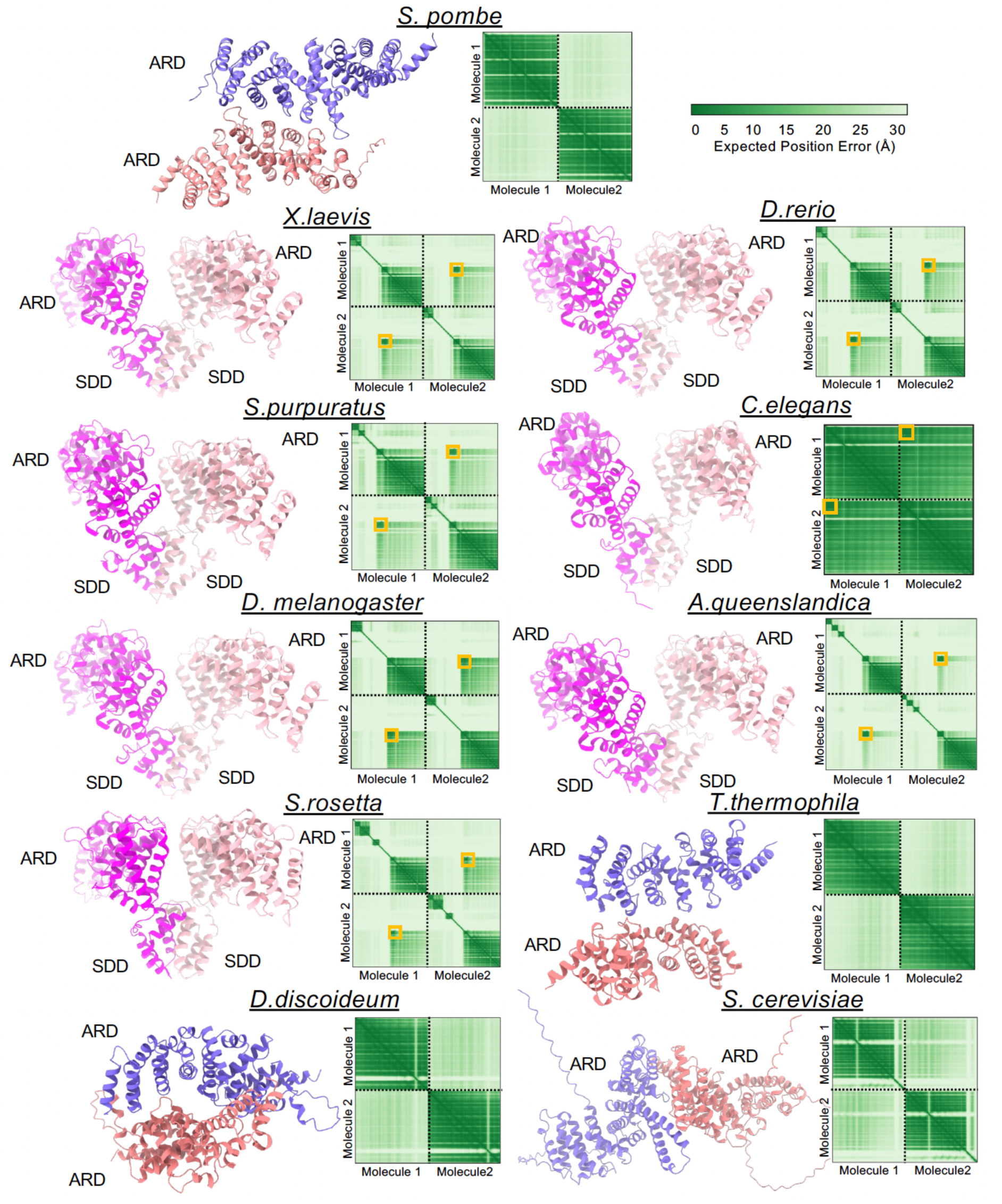
Structural prediction of dimerisation capabilities of WISH/DIP1/SPIN90 family proteins from representative model organisms. For each protein, the left panel shows the predicted AlphaFold model for the SDD and armadillo repeat domains. Subunits in each predicted dimer are coloured diÖerently. The right panel for each protein shows a plot of predicted aligned error (PAE). The SDD domain regions of metazoan with low PAE are highlighted in orange boxes to indicate the confidence of dimer prediction. Note the high PAE between subunits in *S.pombe*, *S. cerevisiae*, *T. thermophila* and *D. discoideum* indicating the low probability of forming a stable dimer.

**Extended Data Table 1.** Cryo-EM data collection, refinement, validation and model building statistics.

|  |  |
| --- | --- |
|  | SPIN90-Arp2/3 complex in the presence of actin (EMDB-52580) (PDB 9I2B) |
| **Data collection and processing** |  |
| Magnification | x81,000 |
| Voltage (kV) | 300 |
| Electron exposure (e-/Å<sup>2</sup>) | 39.2 |
| Defocus range (µm) | -0.9 to -2.4 |
| Pixel size (Å) | 1.06 |
| Symmetry imposed | C2 |
| Initial particle images (no.) | 2,138,010 |
| Final particle images (no.) | 39,104 |
| Map resolution (Å) | 3.0 |
| FSC threshold | 0.143 |
| Map resolution range (Å) | 2.7-4.7 |
| **Refinement** |  |
| Initial model used (PDB code) | 8P94, 6DEE |
| Model resolution (Å) | 3.3 |
| FSC threshold | 0.5 |
| Map sharpening *B* factor (Å<sup>2</sup>) | -62.8 |
| Model composition |  |
| Non-hydrogen atoms | 90,796 |
| Protein residues | 6,128 |
| Ligands | 8 Mg<sup>2+</sup>, 8 ADP, 4 phalloidin |
| *B* factors (Å<sup>2</sup>) |  |
| Protein | 36.44 |
| Ligand | 37.2 |
| R.m.s. deviations |  |
| Bond lengths (Å) | 0.007 |
| Bond angles (°) | 0.839 |
| Validation |  |
| MolProbity score | 1.21 |
| Clashscore | 2.52 |
| Poor rotamers (%) | 0 |
| Ramachandran plot |  |
| Favored (%) | 97 |
| Allowed (%) | 3 |
| Disallowed (%) | 0 |

## REFERENCES

1 Gautreau, A. M., Fregoso, F. E., Simanov, G. & Dominguez, R. Nucleation, stabilization, and disassembly of branched actin networks. Trends Cell Biol 32, 421–432, doi:10.1016/j.tcb.2021.10.006 (2022).

2 Svitkina, T. M. & Borisy, G. G. Arp2/3 complex and actin depolymerizing factor/cofilin in dendritic organization and treadmilling of actin filament array in lamellipodia. J Cell Biol 145, 1009–1026, doi:10.1083/jcb.145.5.1009 (1999).

3 Vinzenz, M. et al. Actin branching in the initiation and maintenance of lamellipodia. J Cell Sci 125, 2775–2785, doi:10.1242/jcs.107623 (2012).

4 Duleh, S. N. & Welch, M. D. WASH and the Arp2/3 complex regulate endosome shape and trafficking. Cytoskeleton (Hoboken) 67, 193–206, doi:10.1002/cm.20437 (2010).

5 Dai, A., Yu, L. & Wang, H. W. WHAMM initiates autolysosome tubulation by promoting actin polymerization on autolysosomes. Nat Commun 10, 3699, doi:10.1038/s41467-019-11694-9 (2019).

6 Machesky, L. M., Atkinson, S. J., Ampe, C., Vandekerckhove, J. & Pollard, T. D. Purification of a cortical complex containing two unconventional actins from Acanthamoeba by affinity chromatography on profilin-agarose. J Cell Biol 127, 107–115, doi:10.1083/jcb.127.1.107 (1994).

7 Welch, M. D., DePace, A. H., Verma, S., Iwamatsu, A. & Mitchison, T. J. The human Arp2/3 complex is composed of evolutionarily conserved subunits and is localized to cellular regions of dynamic actin filament assembly. J Cell Biol 138, 375–384, doi:10.1083/jcb.138.2.375 (1997).

8 Mullins, R. D., Heuser, J. A. & Pollard, T. D. The interaction of Arp2/3 complex with actin: nucleation, high affinity pointed end capping, and formation of branching networks of filaments. Proc Natl Acad Sci U S A 95, 6181–6186, doi:10.1073/pnas.95.11.6181 (1998).

9 Amann, K. J. & Pollard, T. D. The Arp2/3 complex nucleates actin filament branches from the sides of pre-existing filaments. Nat Cell Biol 3, 306–310, doi:10.1038/35060104 (2001).

10 Robinson, R. C. et al. Crystal structure of Arp2/3 complex. Science 294, 1679–1684, doi:10.1126/science.1066333 (2001).

11 Goley, E. D. & Welch, M. D. The ARP2/3 complex: an actin nucleator comes of age. Nat Rev Mol Cell Biol 7, 713–726, doi:10.1038/nrm2026 (2006).

12 Bieling, P. & Rottner, K. From WRC to Arp2/3: Collective molecular mechanisms of branched actin network assembly. Curr Opin Cell Biol 80, 102156, doi:10.1016/j.ceb.2023.102156 (2023).

13 Machesky, L. M. et al. Scar, a WASp-related protein, activates nucleation of actin filaments by the Arp2/3 complex. Proc Natl Acad Sci U S A 96, 3739–3744, doi:10.1073/pnas.96.7.3739 (1999).

14 Blanchoin, L. et al. Direct observation of dendritic actin filament networks nucleated by Arp2/3 complex and WASP/Scar proteins. Nature 404, 1007–1011, doi:10.1038/35010008 (2000).

15 Panchal, S. C., Kaiser, D. A., Torres, E., Pollard, T. D. & Rosen, M. K. A conserved amphipathic helix in WASP/Scar proteins is essential for activation of Arp2/3 complex. Nat Struct Biol 10, 591–598, doi:10.1038/nsb952 (2003).

16 Rottner, K., Hanisch, J. & Campellone, K. G. WASH, WHAMM and JMY: regulation of Arp2/3 complex and beyond. Trends Cell Biol 20, 650–661, doi:10.1016/j.tcb.2010.08.014 (2010).

17 Ti, S. C., Jurgenson, C. T., Nolen, B. J. & Pollard, T. D. Structural and biochemical characterization of two binding sites for nucleation-promoting factor WASp-VCA on Arp2/3 complex. Proc Natl Acad Sci U S A 108, E463–471, doi:10.1073/pnas.1100125108 (2011).

18 Rodnick-Smith, M., Luan, Q., Liu, S. L. & Nolen, B. J. Role and structural mechanism of WASP-triggered conformational changes in branched actin filament nucleation by Arp2/3 complex. Proc Natl Acad Sci U S A 113, E3834-3843, doi:10.1073/pnas.1517798113 (2016).

19 Zimmet, A. et al. Cryo-EM structure of NPF-bound human Arp2/3 complex and activation mechanism. Sci Adv 6, doi:10.1126/sciadv.aaz7651 (2020).

20 Fäßler, F., Dimchev, G., Hodirnau, V. V., Wan, W. & Schur, F. K. M. Cryo-electron tomography structure of Arp2/3 complex in cells reveals new insights into the branch junction. Nat Commun 11, 6437, doi:10.1038/s41467-020-20286-x (2020).

21 Narvaez-Ortiz, H. Y. & Nolen, B. J. Unconcerted conformational changes in Arp2/3 complex integrate multiple activating signals to assemble functional actin networks. Curr Biol 32, 975–987 e976, doi:10.1016/j.cub.2022.01.004 (2022).

22 Chou, S. Z., Chatterjee, M. & Pollard, T. D. Mechanism of actin filament branch formation by Arp2/3 complex revealed by a high-resolution cryo-EM structureof the branch junction. Proc Natl Acad Sci U S A 119, e2206722119, doi:10.1073/pnas.2206722119 (2022).

23 Ding, B. et al. Structure of Arp2/3 complex at a branched actin filament junction resolved by single-particle cryo-electron microscopy. Proc Natl Acad Sci U S A 119, e2202723119, doi:10.1073/pnas.2202723119 (2022).

24 Campellone, K. G., Lebek, N. M. & King, V. L. Branching out in different directions: Emerging cellular functions for the Arp2/3 complex and WASP-family actin nucleation factors. Eur J Cell Biol 102, 151301, doi:10.1016/j.ejcb.2023.151301 (2023).

25 Chavali, S. S. et al. Cryo-EM structures reveal how phosphate release from Arp3 weakens actin filament branches formed by Arp2/3 complex. Nat Commun 15, 2059, doi:10.1038/s41467-024-46179-x (2024).

26 Liu, T. et al. Cortactin stabilizes actin branches by bridging activated Arp2/3 to its nucleated actin filament. Nat Struct Mol Biol, doi:10.1038/s41594-023-01205-2 (2024).

27 Pollard, T. D. & Borisy, G. G. Cellular motility driven by assembly and disassembly of actin filaments. Cell 112, 453–465, doi:10.1016/s0092-8674(03)00120-x (2003).

28 Wagner, A. R., Luan, Q., Liu, S. L. & Nolen, B. J. Dip1 defines a class of Arp2/3 complex activators that function without preformed actin filaments. Curr Biol 23, 1990–1998, doi:10.1016/j.cub.2013.08.029 (2013).

29 Basu, R. & Chang, F. Characterization of dip1p reveals a switch in Arp2/3- dependent actin assembly for fission yeast endocytosis. Curr Biol 21, 905–916, doi:10.1016/j.cub.2011.04.047 (2011).

30 Balzer, C. J. et al. Synergy between Wsp1 and Dip1 may initiate assembly of endocytic actin networks. Elife 9, doi:10.7554/eLife.60419 (2020).

31 Fukuoka, M. et al. A novel neural Wiskott-Aldrich syndrome protein (N-WASP) binding protein, WISH, induces Arp2/3 complex activation independent of Cdc42. J Cell Biol 152, 471–482, doi:10.1083/jcb.152.3.471 (2001).

32 Luan, Q., Liu, S. L., Helgeson, L. A. & Nolen, B. J. Structure of the nucleation-promoting factor SPIN90 bound to the actin filament nucleator Arp2/3 complex. EMBO J 37, doi:10.15252/embj.2018100005 (2018).

33 Kim, D. J. et al. Interaction of SPIN90 with the Arp2/3 complex mediates lamellipodia and actin comet tail formation. J Biol Chem 281, 617–625, doi:10.1074/jbc.M504450200 (2006).

34 Kim, S. H. et al. Interaction of SPIN90 with syndapin is implicated in clathrin-mediated endocytic pathway in fibroblasts. Genes Cells 11, 1197–1211, doi:10.1111/j.1365-2443.2006.01008.x (2006).

35 Cao, L. et al. SPIN90 associates with mDia1 and the Arp2/3 complex to regulate cortical actin organization. Nat Cell Biol 22, 803–814, doi:10.1038/s41556-020-0531-y (2020).

36 Shaaban, M., Chowdhury, S. & Nolen, B. J. Cryo-EM reveals the transition of Arp2/3 complex from inactive to nucleation-competent state. Nature Structural & Molecular Biology 27, 1009–1016, doi:10.1038/s41594-020-0481-x (2020).

37 Forwood, J. K. et al. Quantitative structural analysis of importin-beta flexibility: paradigm for solenoid protein structures. Structure 18, 1171–1183, doi:10.1016/j.str.2010.06.015 (2010).

38 Wen, K. K. & Rubenstein, P. A. Acceleration of yeast actin polymerization by yeast Arp2/3 complex does not require an Arp2/3-activating protein. J Biol Chem 280, 24168–24174, doi:10.1074/jbc.M502024200 (2005).

39 Donnelly, S. K., Weisswange, I., Zettl, M. & Way, M. WIP provides an essential link between Nck and N-WASP during Arp2/3-dependent actin polymerization. Curr Biol 23, 999–1006, doi:10.1016/j.cub.2013.04.051 (2013).

40 Lim, C. S. et al. SPIN90 (SH3 protein interacting with Nck, 90 kDa), an adaptor protein that is developmentally regulated during cardiac myocyte differentiation. J Biol Chem 276, 12871–12878, doi:10.1074/jbc.M009411200 (2001).

41 Wan Mohamad Noor, W. N. I., et al. Small GTPase Cdc42, WASP, and scaffold proteins for higher-order assembly of the F-BAR domain protein. Sci Adv 9, eadf5143, doi:10.1126/sciadv.adf5143 (2023).

42 Cao, L., Ghasemi, F., Way, M., Jegou, A. & Romet-Lemonne, G. Regulation of branched versus linear Arp2/3-generated actin filaments. EMBO J 42, e113008, doi:10.15252/embj.2022113008 (2023).

43 Booth, D. S. & King, N. The history of Salpingoeca rosetta as a model for reconstructing animal origins. Curr Top Dev Biol 147, 73–91, doi:10.1016/bs.ctdb.2022.01.001 (2022).

44 Brunet, T. & King, N. The Origin of Animal Multicellularity and Cell Differentiation. Dev Cell 43, 124–140, doi:10.1016/j.devcel.2017.09.016 (2017).

45. Biomimetic actin cortices shape cell-sized lipid vesicles. bioRxiv 2023.01.15.524117; doi: 10.1101/2023.01.15.524117 (2023).

46 Cao, L., Huang, S., Basant, A., Mladenov, M. & Way, M. CK-666 and CK-869 differentially inhibit Arp2/3 iso-complexes. EMBO Rep 25, 3221-3239, doi: 10.1038/s44319-024-00201-x (2024).

47 Schindelin, J. et al. Fiji: an open-source platform for biological-image analysis. Nat Methods 9, 676-682, doi:10.1038/nmeth.2019 (2012).

48 Punjani, A., Rubinstein, J. L., Fleet, D. J. & Brubaker, M. A. cryoSPARC: algorithms for rapid unsupervised cryo-EM structure determination. Nat Methods 14, 290–296, doi:10.1038/nmeth.4169 (2017).

49 Bepler, T. et al. Positive-unlabeled convolutional neural networks for particle picking in cryo-electron micrographs. Nat Methods 16, 1153–1160, doi:10.1038/s41592-019-0575-8 (2019).

50 Punjani, A. & Fleet, D. J. 3D variability analysis: Resolving continuous flexibility and discrete heterogeneity from single particle cryo-EM. J Struct Biol 213, 107702, doi:10.1016/j.jsb.2021.107702 (2021).

51 Pettersen, E. F. et al. UCSF ChimeraX: Structure visualization for researchers, educators, and developers. Protein Sci 30, 70–82, doi:10.1002/pro.3943 (2021).

52 Croll, T. I. ISOLDE: a physically realistic environment for model building into low-resolution electron-density maps. Acta Crystallogr D Struct Biol 74, 519–530, doi:10.1107/S2059798318002425 (2018).

53 Kidmose, R. T. et al. Namdinator - automatic molecular dynamics flexible fitting of structural models into cryo-EM and crystallography experimental maps. IUCrJ 6, 526–531, doi:10.1107/S2052252519007619 (2019).

54 Emsley, P., Lohkamp, B., Scott, W. G. & Cowtan, K. Features and development of Coot. Acta Crystallogr D Biol Crystallogr 66, 486–501, doi:10.1107/S0907444910007493 (2010).

55 Afonine, P. V. et al. Real-space refinement in PHENIX for cryo-EM and crystallography. Acta Crystallogr D Struct Biol 74, 531–544, doi:10.1107/S2059798318006551 (2018).

56 Blum, M., et al. InterPro: the protein sequence classification resource in 2025. Nucleic Acids Res 53, D444–D456, doi:10.1093/nar/gkae1082 (2025).

57 Dyer, S. C. et al. Ensembl 2025. Nucleic Acids Res 53, D948–D957, doi:10.1093/nar/gkae1071 (2025).

58 Varadi, M. et al. AlphaFold Protein Structure Database: massively expanding the structural coverage of protein-sequence space with high-accuracy models. Nucleic Acids Res 50, D439–D444, doi:10.1093/nar/gkab1061 (2022).

59 Jumper, J. et al. Highly accurate protein structure prediction with AlphaFold. Nature 596, 583–589, doi:10.1038/s41586-021-03819-2 (2021).

60 Abramson, J. et al. Accurate structure prediction of biomolecular interactions with AlphaFold 3. Nature 630, 493–500, doi:10.1038/s41586-024-07487-w (2024).

61 Hedges, S. B. The origin and evolution of model organisms. Nat Rev Genet 3, 838–849, doi:10.1038/nrg929 (2002).

